# Molecular basis of the Druantia anti-phage defense system

**DOI:** 10.64898/2026.05.12.722862

**Authors:** Haidai Hu, Philipp F. Popp, Nicole R. Rutbeek, Victor Klein-Sousa, Blanca Lopéz-Méndez, Aritz Roa-Eguiara, Damien Piel, Tillmann Pape, Nicholas H. Sofos, Ivo A. Hendriks, Jesper V. Olsen, Inga Songailiene, Alexander Harms, Marc Erhardt, Nicholas M.I. Taylor

**Affiliations:** Department of Drug Design and Pharmacology, Faculty of Health and Medical Sciences, University of Copenhagen, Copenhagen, Denmark; Novo Nordisk Foundation Center for Protein Research, Department of Cellular and Molecular Medicine, Faculty of Health and Medical Sciences, University of Copenhagen, Copenhagen, Denmark; Institute of Biology/Molecular Microbiology, Humboldt-Universität zu Berlin, Berlin, Germany; Department of Experimental Medical Science, Lund University, Lund, Sweden; Protein Production and Characterization Facility, Novo Nordisk Foundation Center for Protein Research, Department of Cellular and Molecular Medicine, Faculty of Health and Medical Sciences, University of Copenhagen, 2200 Copenhagen, Denmark; Institute of Food, Nutrition and Health (IFNH), Department of Health Sciences and Technology (D-HEST), ETH Zurich, Zurich, Switzerland; Core Facility for Integrated Bioimaging (CFIB), Center for Core Facilities (CCF), Faculty of Health and Medical Sciences, University of Copenhagen, 2200 Copenhagen N, Denmark; Copenhagen Center for Glycocalyx Research, Department of Cellular and Molecular Medicine, Faculty of Health Sciences, University of Copenhagen, Copenhagen, Denmark; Institute of Biotechnology, Life Sciences Center, Vilnius University, Lithuania; Max Planck Unit for the Science of Pathogens, Berlin, Germany

## Abstract

Eukaryotes and prokaryotes have evolved diverse antiviral immune systems, many containing helicase modules central to defense. Druantia are widespread bacterial anti-phage defense systems, each built around a large helicase domain-containing protein, DruE, paired with variable subunits. Here, we investigate the molecular basis of a minimal two-protein module Druantia system, DruH-E. We demonstrate that DruH-E is sufficient to confer robust anti-phage defense. DruE exists in equilibrium between a monomer and an asymmetric dimer, with dimerization required for in vivo immunity. Cryo-EM structures define DruE asymmetric dimer assembly and its dsDNA unwinding mechanism, revealing a topologically closed architecture that is specifically activated by dsDNA substrates with a 3′ overhang. We further identify DruH as an ssDNA-binding protein regulated by a metabolic switch, where its activity is inhibited by ATP at physiological concentrations through direct competition with ssDNA. Supported by mass spectrometry and single-cell microscopy data, we establish key determinants of the Druantia defense system and reveal how it mediates a direct antiviral immune mechanism.

## Introduction

Bacteria have evolved multiple defense strategies to counter bacteriophage (phage) infection, forming sophisticated bacterial immune systems (Hampton, Watson and Fineran, 2020; Georjon and Bernheim, 2023; Mayo-Muñoz *et al*., 2024; Hochhauser and Sorek, 2025). By sensing phage invasion, bacteria respond with dedicated anti-phage function to either protect individual cells from phage infection or sacrifice infected cells to abort phage replication and maintain bacterial survival at the population level (Aframian and Eldar, 2023). Bioinformatic analyses and experimental validations have greatly expended the pool of bacterial immune systems (Georjon and Bernheim, 2023; Ledvina and Whiteley, 2024). Recent progress in elucidating the mechanisms of newly identified anti-phage systems has begun to reveal diverse bacterial protection strategies. Among them, many systems have helicase domains (Tesson, Rémi Planel, *et al*., 2024).

Druantia represents one of the most widespread anti-phage defense systems (Doron *et al*., 2018). It is primarily found within bacterial defense islands and is detected in approximately 5.4% of known bacterial genomes (Payne *et al*., 2022). Four Druantia subtypes have been initially identified, all built around a shared protein, DruE, which is predicted to be a dsDNA helicase and has been recently regrouped into the YprA (renamed as MrfA; <u>m</u>itomycin repair factor A) helicase family, a representative of the superfamily 2 (SF2) helicase (Bell *et al*., 2025) (**Fig. 1A, B**). The *druE* gene encodes a protein of approximately 2,000 amino acids and exhibits extensive sequence divergence from MrfA (**Fig. 1C**). It has also been reported that homologs of the *druE* gene are present in other defense systems, including ARMADA, among others (Ofir *et al*., 2018; Bell *et al*., 2025). The different Druantia subtypes include variable accessory genes. Type I and type II Druantia systems are multigene operons. In contrast, Druantia III that accounts for 68% of known Druantia operons contains only a single additional gene, *druH*. This gene encodes a protein of approximately 1,100 amino acids with unknown function (**Fig. 1B, C**).

**Figure 1.**
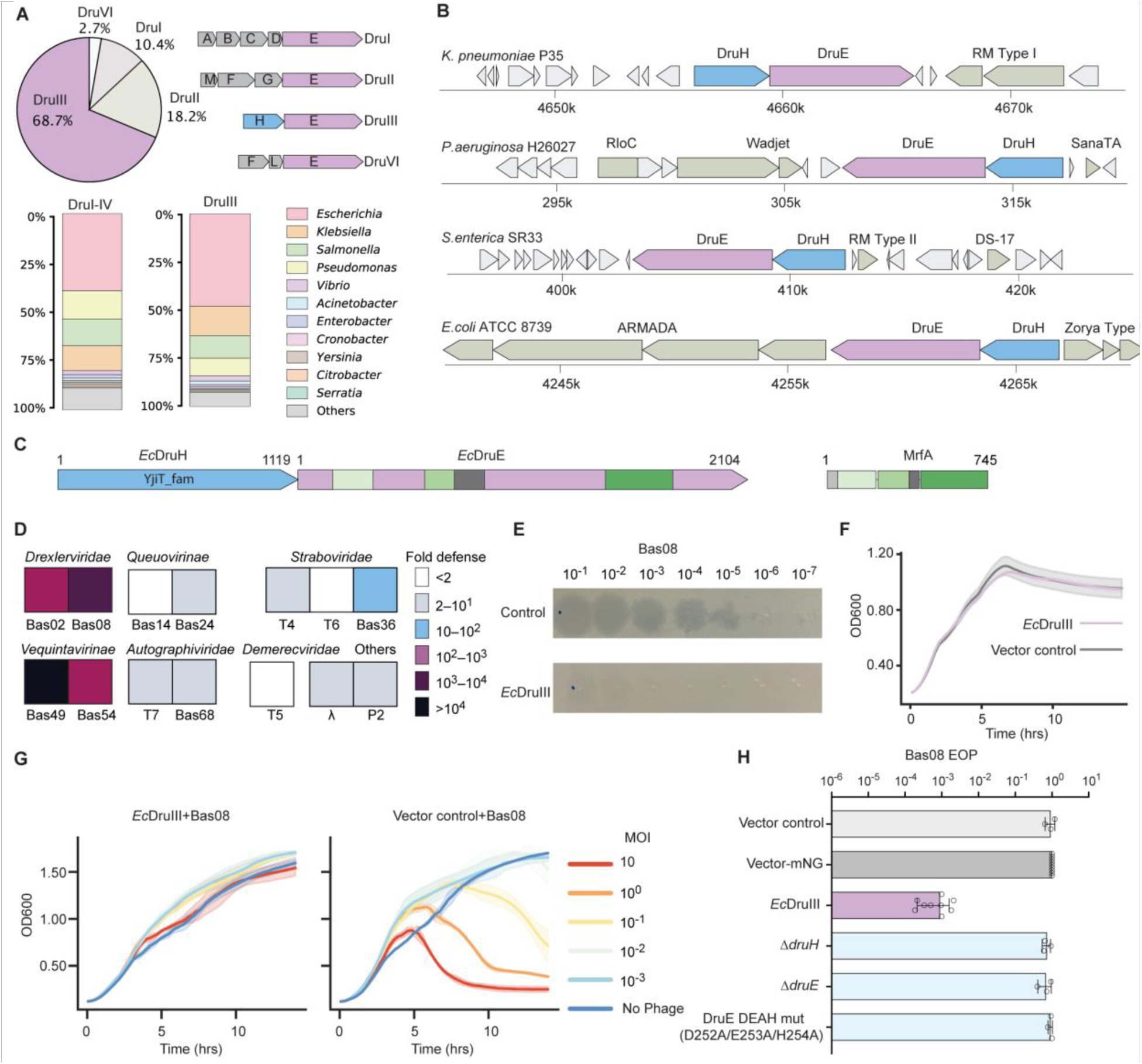
Druantia defense via a direct mechanism. (A) Bacterial genera containing DruIII systems. (B) DruIII operon and gene neighborhoods. (C) Schematic of DruH, DruE and MrfA proteins. (D) *Ec*DruIII defends against diverse *E. coli* phages, measured as fold reduction in plaque formation. (E) Representative plaque assays of phage Bas08 challenging *E. coli* K-12 ΔRM expressing either a vector control or *Ec*DruIII. (F) Growth curve of *E.coli* K-12 ΔRM expressing either vectors control or *Ec*DruIII. (G) Infection time courses of liquid cultures of *E. coli*, with and without *Ec*DruIII, infected at different multiplicities of infection (MOI) of phage Bas08. (H) Effects of mutations on *Ec*DruIII defense against Bas08, measured using EOP assays. Plots or Images in (E), (F) and (G) are representative of at least 3 replicates. Means and standard deviations shown in (D) and (H) are derived from independent biological triplicates.

Here, we investigate the molecular basis of the minimal two-protein module Druantia III anti-phage defense system function using an integrated approach, combining cellular, biochemical, structural, mutagenesis, proteomics and single-cell microscopy analyses. We demonstrate that DruH-E broadly defends against phage infection through a direct immune mechanism. We find that DruE can exist as both a monomer and an asymmetric dimer. We show that the asymmetric dimer, a large ∼500 kDa complex, is necessary for Druantia immunity *in vivo*. Functional *in vitro* reconstitutions show that DruE unwinds dsDNA in a 3′-to-5′ direction and that it requires a 3′ overhang for this activity. This process depends on ATP hydrolysis and is enhanced by substrate-induced asymmetric dimerization. We also discovere that DruH is a DNA-binding protein with a preference for ssDNA, the activity of which is inhibited by ATP at physiological concentrations by competing for the same binding site. Together, by combining structure-guided mutagenesis and functional studies with quantitative mass spectrometry and single-cell microscopy data, our results define key determinants of Druantia-mediated immunity and support a model where DruH and DruE act as a highly coordinated, ATP-regulated complex to detect and directly defend host cells from invading phages.

## Results

### Druantia defends directly against phage infection

We cloned the two-gene Druantia III operon from *E.coli* strain ATCC 8739, together with its native promoter (*Ec*DruIII, a ∼15 kb gene fragment), into a vector (**Fig. S1A**) for expression in *E. coli* K-12 ΔRM, which lacks a native Druantia defense system and additionally all other known restriction modification systems (Maffei *et al*., 2021). To characterize the defense pattern of *Ec*DruIII against phages, we selected representative phages from the BASEL collection (Maffei *et al*., 2021; Humolli *et al*., 2025) as well as several model *E. coli* phages, forming a pool of 14 phages from different families. We then challenged the *Ec*DruIII-containing cells with the selected phage pool and found that *Ec*DruIII provides protection against diverse phages, with the strongest defense observed against phages from the *Drexlerviridae* family and *Vequintavirinae* subfamily (**Fig. 1D, E** and **S1B**). We also observed that Druantia does not affect cell growth in the absence of phages when expressed under the native promoter(**Fig. 1F**). We selected phage Bas08 to assess *Ec*DruIII-mediated protection at the population level in liquid culture. In double-layer agar plaque assays, *Ec*DruIII provides approximately 10^3^-fold protection against Bas08. Upon infection at varying multiplicities of infection (MOIs), *Ec*DruIII-expressing cultures showed no evidence of collapse, and cell growth remained unaffected even at high MOIs **(Fig. 1G)**. These results indicate that *Ec*DruIII protects against phage infection without causing premature cell death. Deletion of either DruH or DruE abolishes defense activity, demonstrating that both are needed for *Ec*DruIII immunity. Furthermore, mutations of the conserved DEAH motif of DruE, predicted to be essential for ATP hydrolysis, lead to a complete loss of defense, confirming that the enzymatic activity of DruE is required for the *Ec*DruIII immunity (**Fig. 1H**).

### DruE exists as a monomer and an asymmetric dimer

To explore the molecular basis of *Ec*DruIII immunity, we initially sought to co-purify DruE and DruH from the plasmid used for in vivo assays (**Fig. S1A**). However, we did not observe complex formation: neither affinity-purified DruE nor affinity-purified DruH pulled down the other protein (**Fig. S1C**). This contrasts with many defense systems in which defense proteins preassemble into stable complexes that are required to mediate anti-phage activity prior to viral infection (Cheng *et al*., 2023; Duncan-Lowey *et al*., 2023; Gao *et al*., 2023; Antine *et al*., 2024; Li *et al*., 2024), suggesting that DruE and DruH may require a phage invasion-derived cue to trigger complex assembly, or that the two proteins may function independently and/or sequentially upon activation. We therefore first focused on the functional and structural characterizations of the individual proteins.

DruE eluted from size-exclusion chromatography (SEC) with a peak position substantially earlier than that expected for a monomer, indicating multimerization. We further validated and quantified this observation using SEC coupled with multi-angle light scattering (SEC-MALS). SEC-MALS analysis indicates that the weight-average molecular mass of purified DruE gradually increases with protein concentration. At concentrations around 10 nM, DruE exists as a monomer and, as the concentration increases, a monomer-dimer equilibrium is observed, with an estimated dissociation constant (*K*_D_) of 0.5 µM (**Fig. 2A-C**). Mass photometry experiments confirmed that the DruE monomer is the main species at low concentrations (∼10 nM) (**Fig. S1D**). To capture the structures of both DruE states, we prepared cryo-EM grids using a high protein concentration of 3.6 µM, at which we expected the dimeric form to be present. Single-particle analysis revealed the existence of both forms and yielded overall resolutions of the DruE monomer and dimer at around 2.7 Å and 3.0 Å, respectively (**Fig. S1E-I, Table S1**). The high-resolution maps were of sufficient quality to build atomic models for most residues in both states.

**Figure 2.**
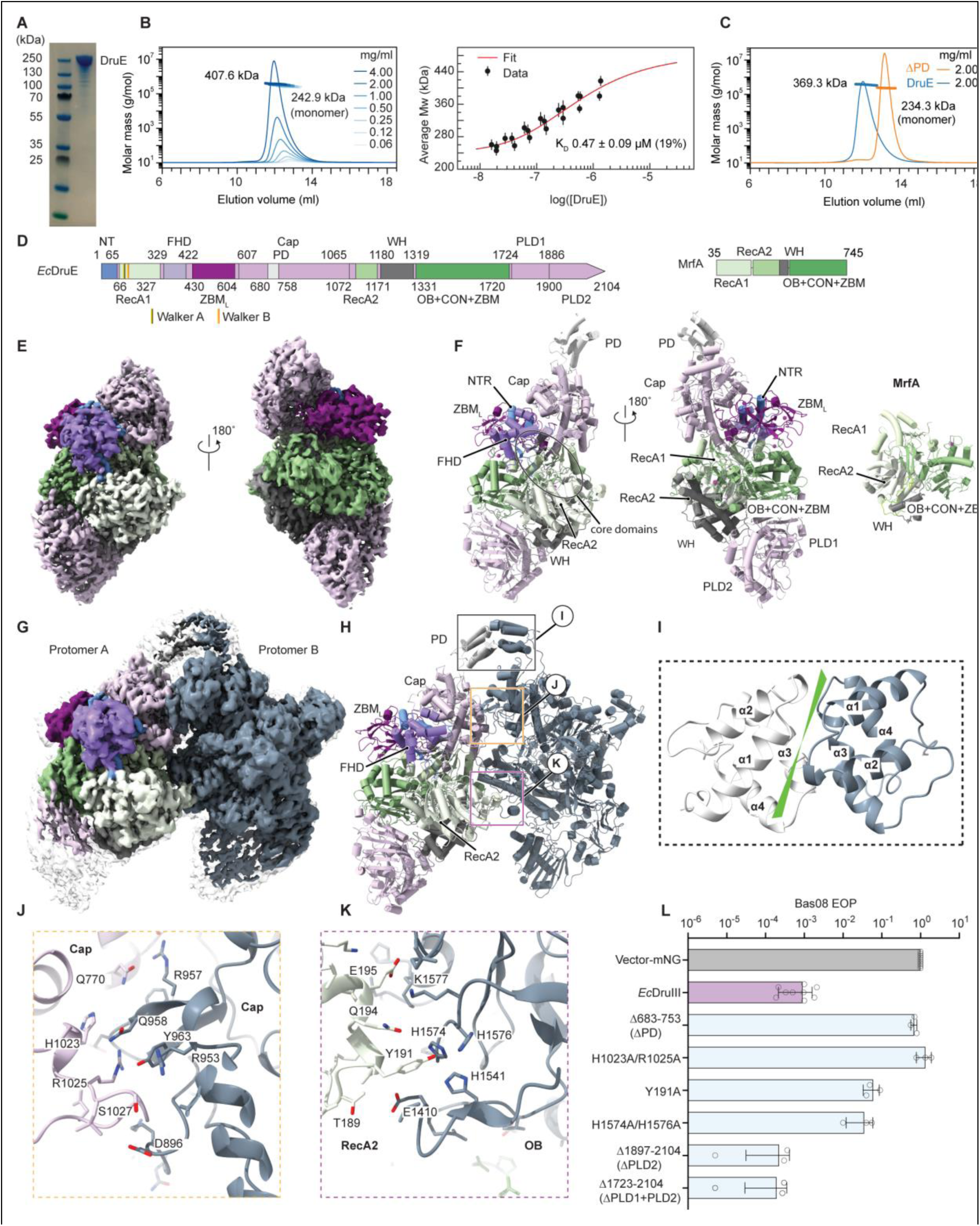
Cryo-EM structures of DruE monomer and asymmetric dimer (A) Representative SDS-PAGE gel of purified DruE. (B) SE C–MALS analysis of DruE at different concentrations, including the derived dimerization constant. (C) SEC–MALS analysis of DruE and the DruE ΔPD mutant (ΔK683–N753). (D) Schematic of DruE and MrfA domains. (E) Cryo-EM map of the DruE monomer. (F) Ribbon representation of monomeric DruE highlighting domains, with comparison to the apo form of MrfA (PDB ID: 6ZNS). (G) Cryo-EM map of the DruE asymmetric dimer. The semi-transparent map represents the Cryo-EM density at a low contour level (σ = 0.015). (H) Ribbon model representation of the DruE asymmetric dimer. (I) Top view of the dimerized DruE PD subdomain and interface I interactions. (J) Close-up view of the interactions of DruE at domain assembly interface II. (K) Close-up view of the interactions at domain assembly interface III. (L) Effects of dimerization interface mutations on *Ec*DruIII defense against Bas08, measured using EOP assays. Images or plots in (A), (B) and (C) are representatives of at least three replicates. Means and standard deviations shown in (L) are derived from independent biological triplicates.

The architecture of the DruE protein displays a core fold resembling that of MrfA (Roske *et al*., 2021), comprising two RecA-like domains (RecA1 and RecA2), an extended antiparallel β-sheet oligonucleotide-binding (OB) fold, a connector (CON) domain that links the OB fold to a zinc-binding module (ZBM) and an atypical ‘winged’ helix-turn-helix domain located at the bottom of the core fold, in which the typical loops or small β-sheet ‘wings’ (Aravind *et al*., 2005) are replaced by three helices (**Fig. 2D-F**). The two highly conserved Walker motifs, which are defining features for all SF2 helicases, are both located within the first RecA domain: the Walker A motif TGSGKT (T117–T122) and the Walker B motif DEAH (D251–H254) (**Fig. 2D**). Besides these shared structural elements, DruE has several unique domains. These domains peripherally surround the core fold and account for approximately 60% of the protein mass, including a compact five-helix domain (FHD) that is spatially bridged by the DruE N-terminal region (NTR) to another large ZBM ( termed as ZBM_L_) that contains two zinc-binding motifs (**Fig. S2A**), as well as a domain that is positioned at the top of the DruE structure and caps the core fold (designated as Cap domain) (**Fig. 2F**). However, in contrast to the previous annotation (Bell *et al*., 2025), the Cap domain has no known homologous structure. Joined by the cap domain, the NTR locks the core fold to form a toroid-shaped topologically closed conformation (**Fig. S2A**). In addition, within the Cap domain, DruE is predicted to contain a four-helix bundle subdomain, which protrudes (termed as PD subdomain; P680-Q758) from the Cap domain. In-gel digestion followed by mass spectrometry of the DruE sample prepared for cryo-EM confirmed the presence of this region during purification (**Fig. S2B**); however, this region appears to be flexible relative to the rest of the structure and does not display well-defined density in the cryo-EM map (**Fig. 2E, F**). Furthermore, DruE contains two phospholipase D (PLD)-like domains at its C terminus, which are located at the lower part of the protein structure. The two PLD-like domains lack the conserved canonical H(X)K(X)₄D catalytic motif required for nuclease activity (Gottlin *et al*., 1998; Yang, 2011) (**Fig. S2C**). Indeed, some DruE proteins from DruIII systems are missing either one or both PLD-like domains (**Fig. S2D**), raising the possibility that they are redundant. Deleting the second PLD-like domain or removing both PLD-like domains does not affect Druantia defense activity, supporting that these two PLD-like domains are dispensable for *Ec*DruIII immunity (**Fig. 2L**).

The DruE dimer accounts for 30% of the entire particle population after the initial round of 3D classification (**Fig. S1G**). Dimerization is mediated by three major interaction interfaces, with the buried area being 2,390 Å², corresponding to approximately 1.4% of the total molecular surface area (163,320 Å²). At the top of the dimerization interface, the PD subdomain of protomer A, which is absent in the monomeric DruE map, becomes ordered and forms a locally 2-fold symmetric interaction with the corresponding part of protomer B (**Fig. 2G, 2H, S3A-C**). This interface I is primarily mediated by the turn between α1 and α2, and α3 helix from each protomer. The PD subdomain is conserved among the DruE of the Druantia III (**Fig. 2I, S3D**). Beneath this dimerized subdomain, the middle region of the Cap domain of protomer A contacts one end of the Cap domain of protomer B, and this interface II is dominated by electrostatic interactions from charged residues (**Fig. 2J**). The last interaction interface (interface III) consists mainly of the DruE core fold. The RecA2 domain, together with the extended OB domain of protomer A, sandwiches the two long helices of the atypical ‘winged’ helix–turn–helix motif of protomer B. In addition, the RecA1 domain of protomer A interacts with the extended OB domain of protomer B, with the interaction mediated by Y191 and three histidine residues (H1415, H1574, and H1576) from the respective protomers (**Fig. 2H, K** and **S3G**). Amongst the interactions, the interfaces II and III break the 2-fold symmetry, which define the structural features of the asymmetric assembly of the DruE dimer (**Fig. 2H, J** and **K**). Superposition of the two protomers within the asymmetric dimer reveals marked conformational differences. Protomer A closely resembles the apo monomeric structure, whereas the protomer B undergoes a substantial conformational change, in which the putative substrate-binding cavity adopts a more open conformation as a consequence of the rearrangement of several long loops and of the outward movement of the Cap domain (**Fig. S3E-F**). This observation suggests that the apo monomer and the protomer A in the asymmetric dimer represent an inactive state, while protomer B, whose conformation is regulated by its partner, likely corresponds to a substrate-ready state.

To explore how dimerization influences DruE function in *Ec*DruIII-mediated anti-phage defense, we designed four mutations targeting the dimer interface, including one truncation in which the protruding four-helix bundle substructure was deleted, and three point mutations at assembly interfaces II and III. All four mutations impaired DruIII defense to varying extents (**Fig. 2L**). Notably, deletion of the PD subdomain (ΔPD; ΔK683-N753) completely abolished anti-phage activity. SEC–MALS analysis showed that this truncation mutant failed to form the asymmetric dimer and existed exclusively as a monomer, indicating that dimerization is crucial for DruE immunity (**Fig. 2C**). Similarly, alanine substitutions of H1023 and R1025 at the interaction interface II (**Fig. 2J**), which predicted to disrupt Cap domain interactions, also completely eliminated DruIII defense activity. By contrast, alanine substitutions of Y191 or the interacting histidine residues (H1574A and H1576A) at the interface III resulted in a partial reduction of defense activity (**Fig. 2K, L**). Consistent with this observation, the purified Y191A mutant retained the ability to form the dimer (**Fig. S3H**). Together, these results reveal that DruE apo states equilibrate between a monomer and dimer, and define the molecular basis of DruE asymmetric dimerization, which is crucial for *Ec*DruIII mediated anti-phage defense.

### DruE dsDNA unwinding mechanism

Structural analysis of apo DruE reveals that it contains an ATP-dependent helicase core domain conserved within the MrfA helicase family. To directly validate the putative DNA helicase activity of DruE and define its substrate requirements, we performed DNA unwinding assays by incubating purified DruE with dsDNA substrates of varying lengths and different single-stranded overhang structures. DruE efficiently unwinds dsDNA substrates containing a 3′ ssDNA overhang or a forked structure. In contrast, it is unable to unwind blunt-ended dsDNA or dsDNA substrates with a 5′ ssDNA overhang (**Fig. 3A** and **Table S2**). Additionally, dsDNA unwinding strictly requires ATP hydrolysis. Replacing ATP with the non-hydrolysable ATP analogue AMP-PNP (adenylyl-imidodiphosphate), a competitive inhibitor of ATP-dependent enzymes, completely abolishes DruE helicase activity (**Fig. 3A**). Consistent with these findings, a purified DruE variant carrying inactivating mutations in the Walker B motif (DruE-D251A/E252A/H254A) renders DruIII-containing cells defenseless against phage infection and exhibits no detectable dsDNA helicase activity (**Fig. 1H, 3B**). The PD subdomain mutant that completely disrupts DruE dimer formation shows reduced, but not abolished, DNA unwinding activity. By contrast, the Y191A mutant, which shows partially reduced defense activity, retains *in vitro* DNA unwinding ability (**Fig. 3B**). SF2 helicases couple ATP hydrolysis to separate dsDNA and move along ssDNA. We assessed DruE ATPase activity using a malachite green assay and found that its ATP hydrolysis is strongly stimulated in the presence of ssDNA (**Fig. S4A**). Mass photometry analysis further showed that at low protein concentrations, ssDNA promotes DruE dimerization, suggesting that ssDNA not only stimulates DruE ATPase activity but also modulates the conformational state of the monomer that facilitates the formation of DruE dimer (**Fig. 3C**). Together, these results demonstrate that DruE is an ATP-dependent dsDNA helicase that specifically requires substrates containing a 3′ ssDNA overhang, and that DruE dimer formation is important for its *in vitro* helicase activity.

**Figure 3.**
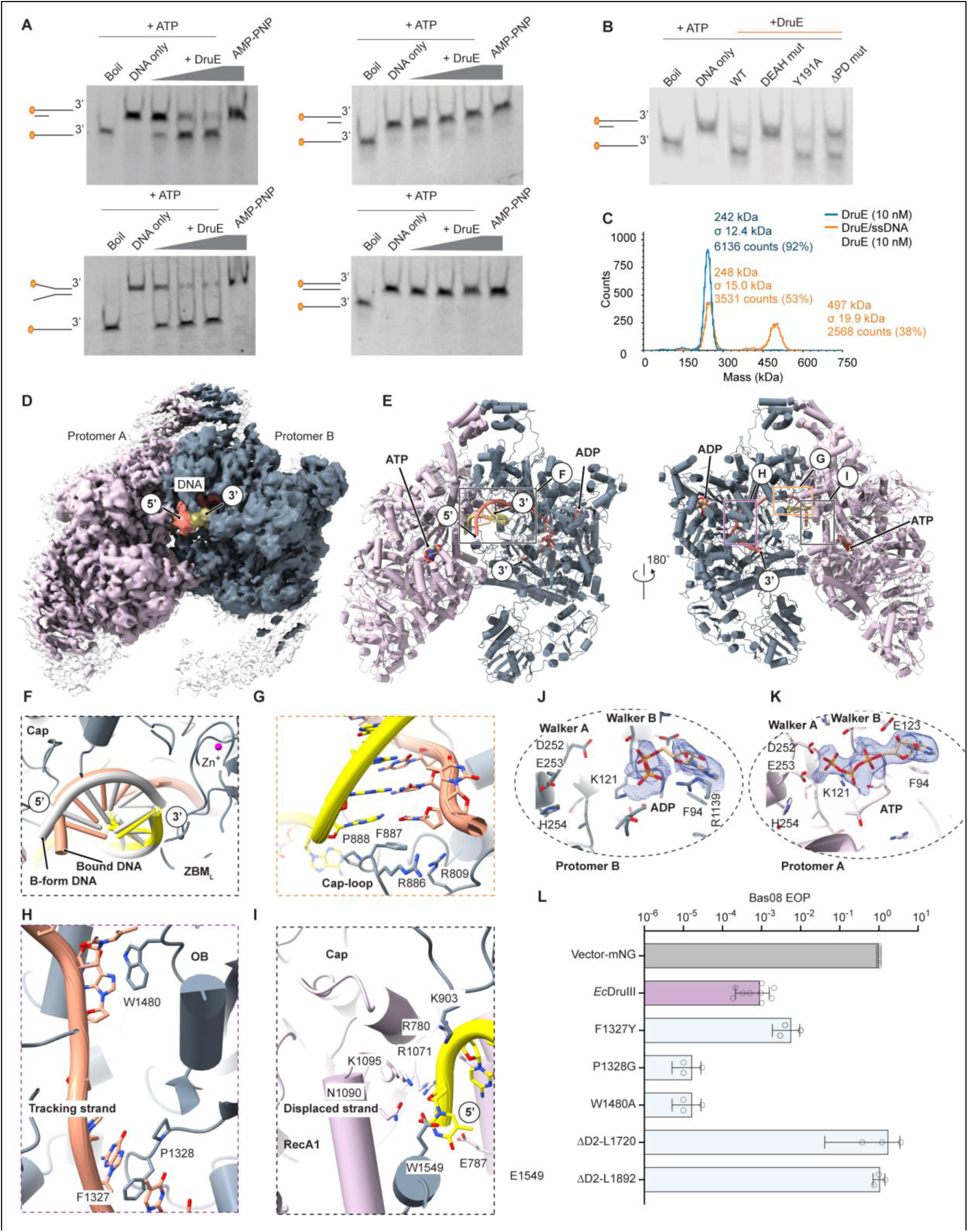
DruE dsDNA unwinding mechanism (A) DruE dsDNA unwinding assays. (B) dsDNA unwinding assays of DruE and various mutants. (C) Mass photometry analysis of DruE alone and DruE bound to ssDNA. (D) Cryo-EM map of the DruE asymmetric dimer with partially unwound dsDNA. The semi-transparent map represents the Cryo-EM density at a low contour level (σ = 0.015) (E) Ribbon model representation of the DruE asymmetric dimer with partially unwound dsDNA. (F) Close-up view of DruE interactions with the dsDNA substrate. An ideal B-form dsDNA (gray) is superimposed. (G) Detailed view of interactions of DruE with dsDNA substrate at the DNA unwinding interface. (H) Detailed view of interactions of DruE with the DNA tracking strand. (I) Detailed view of interactions of DruE with the DNA displaced strand. (J) Close-up view of the ADP molecule identified in protomer B, with EM density overlayed. (K) Close-up view of the ATP molecule identified in protomer A, with EM density overlayed. (L) Effects of mutations in residues interacting with the DNA tracking strand on *Ec*DruIII defense against Bas08, measured by EOP assay. Images or plot in (A), (B) and (C) are representatives of at least 3 replicates. Means and standard deviations shown in (L) are derived from independent biological triplicates.

To define the structural basis of the DruE dsDNA unwinding mechanism, we incubated DruE with the forked dsDNA substrate in the presence of ATP and determined the cryo-EM structure of the asymmetric DruE dimer bound to a partially unwound dsDNA substrate to a nominal resolution of 2.9 Å (**Fig. 3D, E** and **Fig. S4C-D, Table S1**). Remarkably, in this structure, only one DruE protomer (protomer B) makes the primary contribution to dsDNA substrate binding. The cap domain and ZBM_L_ of protomer B engage with the duplex DNA (**Fig. 3E, F)**. These two domains compress and twist the canonical B-form dsDNA, generating an over-twisted supercoiling dsDNA that likely decreases base-pair stability and accumulates repulsive energy to facilitate strand separation (**Fig. 3F, S4E)**. A loop motif from the Cap domain functions as a wedge that reaches beneath the dsDNA separation interface (**Fig. 3E, G)**. Within this loop, residue F887 forms π–π interactions with the last paired bases, and presumably destabilizes base stacking from the neighboring bases. Residue P888, located on one side of F887, pushes apart the first unpaired base on the displaced strand; while residues R886 and R809, located on the other side of F887, twist the tracking strand by interacting with its backbone, resulting in strand separation (**Fig. 3G)**. The tracking strand penetrates deeply into protomer B and its 3′ tail traverses through the toroid-shaped core of DruE, and via the DNA backbone forms extensive electrotactic interactions, confirming that the core fold primarily mediates DNA tracking strand translocation in a 3′-to-5′ direction (**Fig. 3E, H, S4E**). Additional interactions with this DNA tracking strand are provided by F1327 and P1328 from a loop immediately following the atypical ‘winged’ helix–turn–helix motif, as well as a tyrosine residue (W1480) from the OB domain (**Fig. 3H)**; all of these residues that interact with the DNA bases are conserved among DruE (**Fig. S4F-H)**. We hypothesized that the rigidity and the size of these residues would modulate the translocation efficiency of the tracking strand and therefore impact Druantia immunity. To test this hypothesis, we first mutated F1327 at the loop position to tyrosine to increase residue size. The F1327Y mutation showed a decrease in anti-phage defense activity. In contrast, the P1328G mutation, which was intended to increase loop flexibility, unexpectedly displayed an approximately 100-fold increase in anti-phage defense compared to the wildtype. Similarly, replacement of tryptophan with alanine in the OB domain also resulted in an approximately 100-fold increase in anti-phage defense (**Fig. 3L)**. Therefore, our structural analysis and functional data demonstrate that mutations of residues in the core fold responsible for interactions with the nucleobases of the tracking strand finetune *Ec*DruIII anti-phage defense capacity.

Only three unpaired nucleotides of the displaced strand are observed, in contrast to the tracking strand. The last two nucleotides make prominent interactions with the residues from the Cap domain and the RecA1 domain of the opposite promoter (protomer A) (**Fig. 3I)**. Because the forked dsDNA substrate used for structure determination has equal-length arms (**Fig. S4D**), the remaining nucleotides of the displaced strand are likely flexible and disordered. Indeed, the core fold of protomer A adopts an occluded conformation due to the conformational change, leading the displaced strand to pass through a channel framed by the two protomers rather than through the core fold (**Fig. 3I**), the open state of which only accommodates ssDNA in the 3′to 5′ direction as observed in promoter B (**Fig. 3H, S4I-J)**. Notably, density corresponding to a hydrolyzed ATP molecule was identified near the Walker A and Walker B motifs in protomer B, which engages the 3′ tail of the tracking strand, likely an ADP molecule (**Fig. 3J**), suggesting that energy derived from ATP hydrolysis is primarily used to drive translocation of the unwound 3′ ssDNA. By contrast, protomer A contains an intact ATP molecule in the equivalent binding pocket (**Fig. 3K**). These observations explain why the presence of a 3′ ssDNA overhang is required to stimulate DruE helicase activity and support the model that, within the asymmetric DruE dimer, only protomer B actively catalyzes ATP hydrolysis and dsDNA substrate unwinding, while the accompanying protomer functions primarily to maintain the protomer B in a substrate activated conformation and to accommodate the displaced strand. Consequently, strand separation proceeds processivity through ATP hydrolysis only in a single active protomer, consistent with the structural asymmetry observed in the DruE dimer. Combined, these results support that intrinsic helicase activity of DruE is essential for *Ec*DruIII defense and define the dsDNA unwinding mechanism mediated by the DruE asymmetric dimer.

### DruH binds to ssDNA

We next sought to explore the function of DruH and to understand how the DruE dsDNA unwinding mechanism coordinates with DruH to control anti-phage defense. DruE exhibits a deep evolutionary relationship with MrfA, which functions together with MrfB, a 3′ to 5′ DNA exonuclease, in nucleotide excision repair during DNA damage (Burby *et al*., 2018; Manthei *et al*., 2024). As DruH strictly pairs with DruE in almost all sequenced bacterial genomes (**Fig. S5A**), and we did not observe direct interaction between DruH and DruE during purification, or even in the presence of the DruE with its unwound dsDNA substrate (**Fig. S5B-F**), these naturally led us to hypothesize that DruH engages with the ssDNA product generated by DruE during its unwinding process. However, DruH contains no predicted nuclease domain or characteristic motifs related to known DNA binding protein families or folds. We purified DruH and observed that, when incubated with either the dsDNA substrate used to characterize DruE activity or the ssDNA product generated by DruE unwinding, DruH displayed DNA-binding activity (**Fig. 4A, S5G**) with a clear preference for ssDNA over blunt dsDNA (**Fig. 4A**). This binding was enhanced by divalent metal ions, particularly Zn²⁺ (**Fig. 4B, S5H**). However, no DruH nuclease activity was detected under any tested conditions (**Fig. S5I, S7A**).

**Figure 4.**
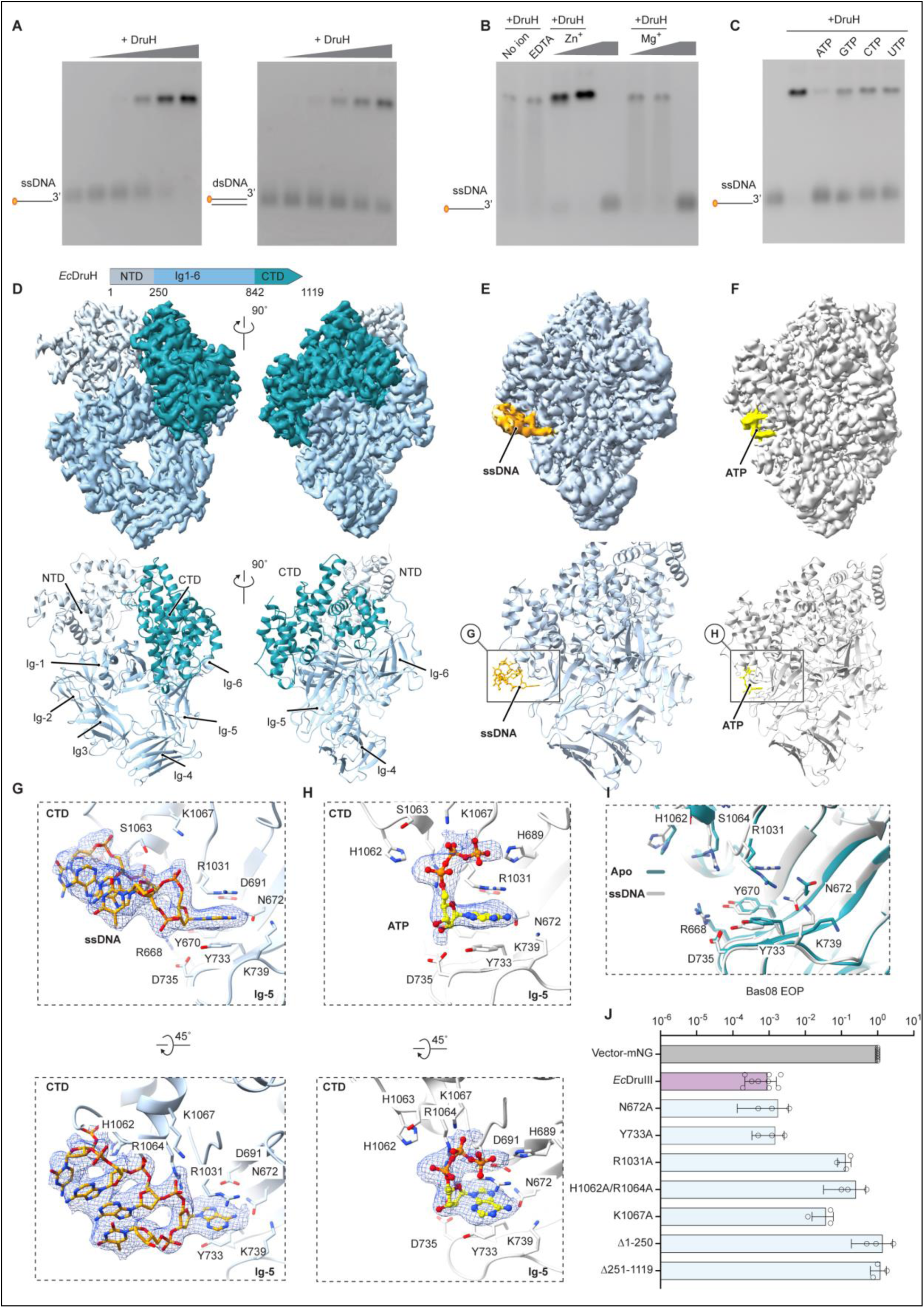
DruH is a ssDNA binding protein (A) DruH DNA binding assays. (B) DruH ssDNA binding in the presence or absence of ions. (C) DruH ssDNA binding in the presence of 1mM NTPs, see Materials and Methods. (D) Cryo-EM map and model of apo DruH. (E) Cryo-EM map and model of DruH in complex with ssDNA. (F) Cryo-EM map and model of DruH in complex with ATP. (G) Detailed view of DruH interactions with ssDNA. (H) Detailed view of DruH interactions with ATP. (I) Close-up view of conformational changes in residues near the substrate-binding site. (J) Effects of mutations in DruH residues interacting with ssDNA on *Ec*DruIII defense against Bas08, measured by EOP assays. Images in (A), (B) and (C) are representatives of at least 3 replicates. Means and standard deviations shown in (J) are derived from independent biological triplicates.

### Structural basis of DruH ssDNA binding and inhibition by ATP

To elucidate the DruH ssDNA binding site, we first determined the cryo-EM structure of DruH in its apo form at 2.2 Å resolution, with a high-quality electrostatic potential map for nearly the entire protein (**Fig. 4D, S6A, Table S1**). DruH comprises a globular N-terminal domain connected via six immunoglobulin-like subdomains (Ig1-6) to its C-terminal domain, which in turn interacts with the N-terminal domain (**Fig. 4D**). The overall structure of DruH is in good agreement with the AlphaFold-predicted model, except for the N-terminal globular domain, which is predicted to adopt multiple conformations relative to the rest of the protein (**Fig. S6B**). Deletion of either the N-terminal globular domain or the C-terminal remainder of DruH resulted in loss of Druantia defense activity *in vivo* (**Fig. 4J**). The electrostatic surface of DruH is largely negatively charged, precluding confident prediction of the DNA-binding site based on the apo structure (**Fig. S6C**).

We then incubated DruH with ssDNA and determined cryo-EM structure of this complex. The 3D reconstruction of the complex, resolved at 2.3 Å resolution, clearly reveals density signal corresponding to ssDNA (**Fig. 4E, S6D, Table S1**). The ssDNA-binding site is located at the interface between the fifth immunoglobulin-like subdomain and the C-terminal domain of DruH, with no other regions contributing to ssDNA binding (**Fig. 4E, G**). At the DruH ssDNA binding interface, one ssDNA nucleobase inserts into a cavity surrounded by several well-conserved residues. DruH R1031 and Y733 sandwich the nucleobase through π-cation interaction and π-π stacking. The aromatic ring of Y670 further stabilizes the interaction by providing lateral support, while deep into the cavity, N672 makes a hydrogen bond with the nucleobase. In addition, positively charged residues K1067, H1062 and R1064 coordinate the phosphate group of the ssDNA backbone from the 5′ end of the inserted nucleobase (**Fig. 4G**). We were only able to confidently model five nucleotides (of the 59-nt substrate); the remaining nucleotides are disordered and exhibit cloud-like electrostatic potential in the cryo-EM map, suggesting that they are conformationally dynamic. Comparison of the apo DruH structure with the DruH ssDNA complex shows that only residues surrounding the DNA-binding motif undergo side chain rearrangements to accommodate ssDNA binding. Particularly R1031, whose side chain entirely occupies the nucleobase-inserted cavity in the DruH apo form, rearranges upon ssDNA binding to expose this unique cavity in the complex responsible for the nucleobase binding (**Fig. 4I**). No major global conformational change was detected between the DruH apo and ssDNA-bound states (**S6E-G**). These findings support that DruH is an ssDNA-binding protein and likely to bind the ssDNA product generated by DruE dsDNA unwinding during Druantia function. Consistent with the structural observations, substitution of R1031 with alanine, results in an approximately 90% reduction in *Ec*DruIII mediated phage defense, whereas mutations of H1062 and R1064 lead to a similar reduction in protection. By contrast, individual alanine mutations of N672 or Y733 have minor effects on *Ec*DruIII defense (**Fig. 4J**). Together, these results establish DruH as an ssDNA-binding protein and reveal critical molecular determinants of the DruH ssDNA interaction that are necessary to support *Ec*DruIII mediated anti-phage defense.

While testing the optimal substrate binding conditions of DruH, we serendipitously found that ssDNA binding by DruH is inhibited by ATP and, to a lesser extent, by other cellular NTPs (**Fig. 4C, S7A**, **B**). DruH does not exhibit ATPase activity, even in the presence of a DNA substrate (**Fig. S4A**). To explore whether ATP inhibition is an allosteric effect resulting from changes in the protein conformational or occurs via a direct inhibitory mechanism, we incubated DruH with both a DNA substrate and ATP and resolved the cryo-EM structure. We were only able to obtain a DruH:ATP complex structure at 2.4 Å resolution (**Fig. 4F, S7C, Table S1**). The structure reveals that the adenine ring of the ATP molecule specifically binds to the ssDNA nucleobase binding cavity. The three phosphate groups of ATP are coordinated by charged residues, including H1062, K1067 and H689. These data support a model where ATP directly inhibits ssDNA binding by competing for the same binding site (**Fig. 4F, H, S7D-F**). Due to this inhibitory effect, physiological *E.coli* cellular ATP levels ( above 3 mM) (Buckstein, He and Rubin, 2008) would likely maintain DruH in an inactive state. However, we note that the observed *in vitro* competition might not fully reflect the inhibitory effect of ATP on DruH ssDNA binding *in vivo*.

### Druantia defends against phage infection at a single cell level

Guided by these structural and functional insights on DruE and DruH, we next investigated how Druantia provides direct anti-phage defense at the cellular level. Since the defensive function of DruE depends on its asymmetric dimer and the expression level of many defense systems correlates with their defensive capacity (Aframian *et al*., 2025), we first measured the copy number of DruH and DruE at the single-level to quantify their abundances under the native *Ec*DruIII protomer. Quantitative mass spectrometry analysis, using *E. coli* protein copy numbers as an external “proteomic ruler” (Schmidt *et al*., 2016) indicated approximately 190 copies of DruH and 470 copies of DruE per cell (**Fig. 5A** and **Table S3)**. Combining these numbers with the estimated *E. coli* cell volume (Volkmer and Heinemann, 2011), we calculated the intracellular concentrations of DruH and DruE to be approximately 0.08 µM and 0.2 µM, respectively. These values suggest that, under the tested conditions, majority of DruE likely exist as monomers (dimerization *K*_D_ ∼0.5 µM), with an approximate DruH:DruE protein stoichiometry of 1:2.

**Figure 5.**
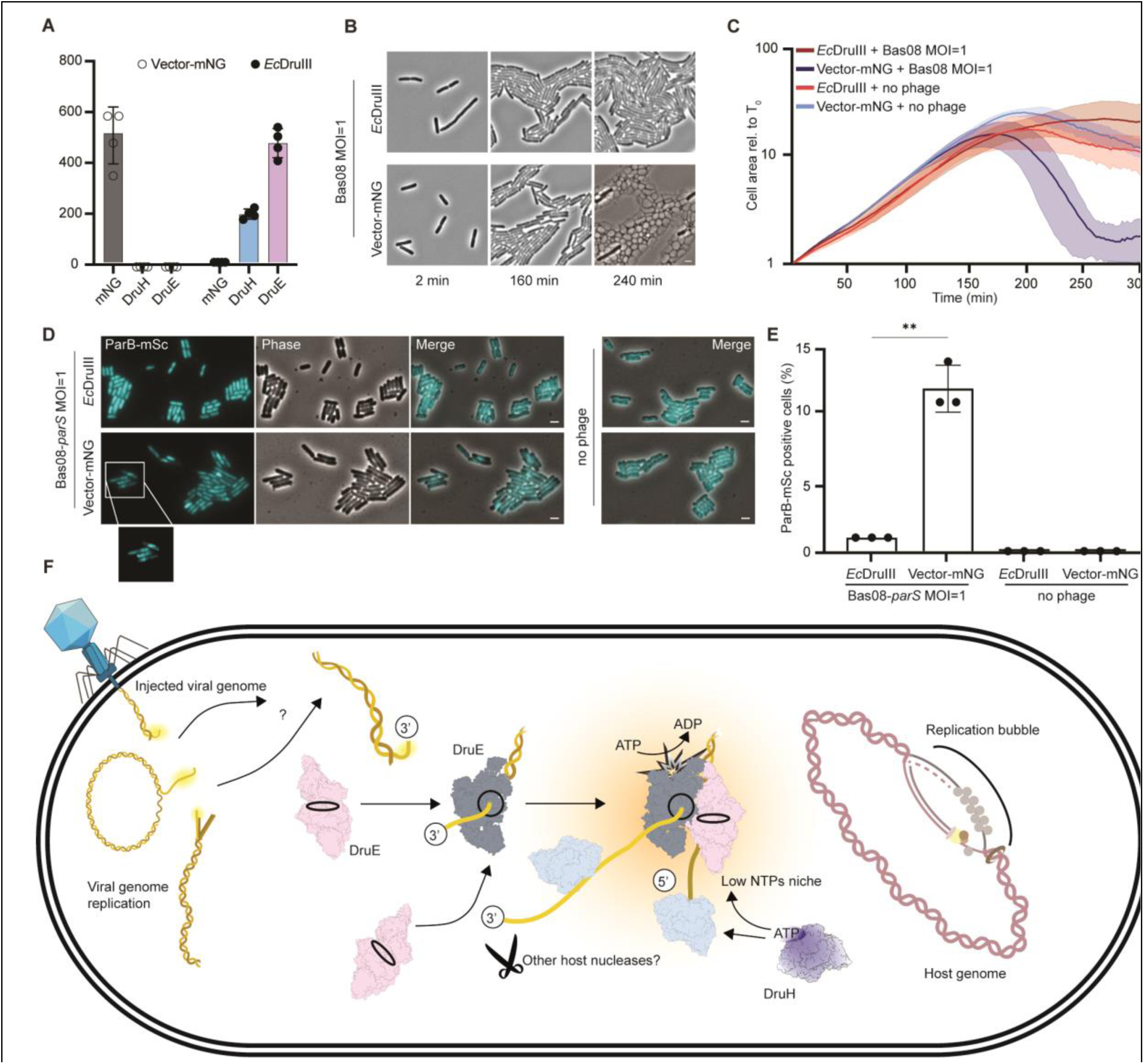
Druantia provides direct anti-phage defense at the single-cell level with a proposed model. (A) Estimated intracellular copy numbers of DruH and DruE in *E. coli* expressing *Ec*DruIII. (B) Representative single-cell time-lapse microscopy images of *E. coli* expressing *Ec*DruIII or an empty vector (vector-mNG) following infection with phage Bas08 at MOI=0.1. Time after infection is indicated. Scale bar is set to 10 µm. (C) Quantification of cell growth during infection measured as cell area relative to the initial time point (T_0_). Shaded areas indicate standard deviation. (D) Visualization of phage DNA during infection using a ParB–*parS* labeling system. Infection assays were performed in an *E. coli* strain constitutively expressing ParB^P1^–mScarlet-I with Bas08-MCP-*parS* at an MOI=1. Representative phase-contrast and fluorescence images at 120 min post-infection show ParB foci in cells carrying the empty vector, indicating the presence of phage DNA. Few foci are observed in *Ec*DruIII-expressing cells or no foci are observed in non-infected controls. Scale bar is set to 2 µm. (E) Quantification of infected cells based on ParB foci formation. (F) Proposed *Ec*DruIII defense model. DruE recognizes 3′ free ends of phage genomic DNA upon infection or during replication. Recognition of 3′ free ends induces conformational changes in DruE, promoting its asymmetric dimerization and unwinding of phage genomic DNA. Phage infection or DruE ATPase activity consume cellular NTPs, which probably relieves the inhibitory effect on DruH. DruH subsequently binds ssDNA generated by DruE, resulting in immobilized phage ssDNA. This could interfere with phage replication and allow for other nucleases to be recruited to degrade phage genomic DNA. The circular replication of *E. coli* would not trigger DruE loading and activation under normal growth conditions.

Type I and type II Druantia contain methyltransferase accessory modules. A recent finding indicates that type I Druantia modifies the host genome to allow to selectively recognize invading phage genomic DNA(Ojima *et al*., 2024). We next examined whether Druantia III also modifies host genomic DNA. To test this, we performed nanopore sequencing of the bacterial genome in the presence or absence of *Ec*DruIII. Analysis of the bacterial genomic DNA did not reveal increased methylation at the global level, ruling out the possibility that Druantia III distinguishes self from non-self DNA by introducing modifications to the host genome (**Fig. S8A-D**). This is consistent with the structural analysis that DruE and DruH proteins contain no methyltransferase domains.

We then explored Druantia defense at the single-cell level. Using single-cell time-lapse microscopy, we observed that *Ec*DruIII-expressing cells infected with Bas08 (MOI=1) continued to grow, whereas the majority of the bacterial population carrying the empty vector began to collapse at around 200 minutes and was cleared by 300 minutes after phage infection. These observations further corroborate that *Ec*DruIII mediates direct defense by protecting individual infected cells (**Fig. 5B, C**). To directly monitor phage DNA during infection, we genetically engineered Bas08 to carry a *parS* sequence inserted downstream of the MCP gene in the phage genome (see Methods). Using this modified phage, we performed time-lapse microscopy infection assays at an MOI=0.1 in an *E. coli* strain constitutively expressing ParB^P1^ translationally fused to mScarlet-I from the chromosome (Li and Austin, 2002). After Bas08 infection, ParB foci, were detected in roughly half of the cells carrying the empty vector at around 200 minutes, indicating the presence of double-stranded phage genomic DNA inside the host. In contrast, no foci formation was observed in cells expressing *Ec*DruIII or in non-infected controls (**Videos S1** and **Videos S2**). We confirmed this trend by running the infection assay with an increased MOI=1 in liquid and visualized single-cells using microscopy after 120 minutes post infection (**Fig. 5D, E**). Since *Ec*DruIII does not have active nuclease domains, we postulate that the lack of ParB foci is caused by DruH binding to unwound phage ssDNA produced by DruE. This would prevent ParB binding which requires a dsDNA *parS* site (Taylor *et al*., 2015; Couturier *et al*., 2023). Together, these single-cell *in vivo* data support a model in which Druantia directly protects cells from phage infection by interfering with phage genome DNA.

## Discussion

Our study provides key molecular insights into the anti-phage defense activity mediated by the minimal two-protein module Druantia system, which exploits a direct antiviral mechanism through the coordinated action of DruE and DruH. We demonstrate that DruE functions as dsDNA helicase requiring a specific substrate structure to unwind dsDNA in a 3′-to-5′ direction, whereas DruH is a ssDNA binding protein whose activity is inhibited by NTPs, most notably ATP. Our findings support a mechanistic anti-phage defense model, in which upon phage infection, DruE recognizes free 3′ ssDNA of phage genomic DNA and is stimulated to assemble into an asymmetric dimer that facilitates unwinding of the phage genome (**Fig. 5F**). Indeed, 3′ overhangs are common DNA structural features in the life cycle of many phages, such as 3′ cohesive ends during genome injection and rolling circle replication or 3′ terminal overhang repeats during genome replication, which are used for concatemer formation in many dsDNA phages (Hoelz, Hickey and Malkas, 2004; Casjens and Gilcrease, 2009; Molineux and Panja, 2013). In the meantime, DruH is activated and binds to the ssDNA generated by DruE to prevent separated DNA strands from reannealing, thereby immobilizing the phage genome. Importantly, unlike most helicases, DruE forms a topologically closed, toroid-shaped structure. This architecture means that activation or loading of DruE onto a DNA substrate requires linear 3’ overhang of ssDNA to initiate helicase activity. Consequently, under normal growth conditions, the circular replication of *E. coli* genomic DNA, even with multiforked chromosomes (Youngren *et al*., 2014), would not trigger DruE loading and activation, in this way ensuring a direct defense mechanism, which responds selectively to invading phage DNA (**Fig. 5F**).

DruE is a key member of the MrfA helicase family, which is widespread in many anti-phage defense systems and shows great sequence diversity. The functional and high-resolution structural insights presented in this study provide a foundation for understanding DruE helicase in Druantia defense systems. Notably, the tertiary structure of the DruE dimer bound to partially unwound DNA reveals how one DruE subunit recognizes the specific dsDNA structure to promote ATP hydrolysis and ssDNA strand translocation, while the other subunit captures the displaced strand, highlighting the structural advantage of this asymmetric assembly. However, the protruding domain required for DruE dimer assembly and anti-phage defense seems to be specifically conserved only within Druantia III systems (**Fig. S3D**). Other Druantia subtypes lack this subdomain, raising the question of whether alternative structural elements in DruE from these systems perform a similar function, or whether other Druantia subtypes exploit divergent defense mechanisms by coupling DruE with diverse accessory proteins. The gain-of-function phenotype of two DruE mutants are particularly interesting, as the mutated residues in DruE interact with the nucleobase of the tracking strand. Phages with extensive modifications, such as T4, are likely to stall the helicase activity by clashing with these residues and thereby suppress Druantia immunity. Indeed, DruE provides only a minor defense against wildtype T4. However, it provides strong defense activity against T4C, in which the cytosine modification was removed (Wang *et al*., 2023).

We further provide *in vitro* evidence that DruH ssDNA binding activity is inhibited by ATP at cellular concentrations, which would maintain DruH in an inactive state under normal growth conditions *in vivo*. Manipulation of the free nucleotide pool has been shown to be a successful antiviral strategy employed by host immune systems from prokaryotes to eukaryotes. Phage infection may lead to massive consummation of cellular NTPs as a result of rapid phage genome transcription and replication. Sensing the drop of the cellular NTPs during viral infection can trigger the activation of many bacterial defense systems, such Gabija (Cheng *et al*., 2023), KELShedu (Zhang *et al*., 2025) and Ppl defense system (Xu *et al*., 2026). We propose that phage infection may relieve NTP-mediated inhibition of DruH, particularly the inhibition mediated by ATP. Additionally, ATP consumption during DruE-mediated dsDNA unwinding could generate a localized low-ATP microenvironment, promoting dissociation of ATP from DruH in close proximity to the DruE activation site, where DruH binds and immobilizes ssDNA produced by DruE. Alternatively, the produced ssDNA could further recruit and bind the apo monomeric DruE, stimulating its ATPase activity, increasing the local ATP depletion rate.

Immobilization of the phage genome without degrading it would create time for other host defense systems to act, as the Druantia III defense system lacks an apparent nuclease module. Each bacterial genome encodes on average five distinct anti-phage defense systems, which are presumed to function in a layered mode to protect the cell against phage infection, such that upon failure of one defense mechanism, others can provide backup (Tesson and Bernheim, 2023; Tesson, Remi Planel, *et al*., 2024). Druantia III system has been observed to exhibit synergistic effects when paired together with other defense systems, including type II Zorya systems (Wu *et al*., 2024; Li *et al*., 2025). Indeed, the defense island from which we cloned *Ec*DruIII contains a Zorya II system (**Fig. 1A**). The Zorya system is proposed to function at early stage of phage infection by detecting an invasion signal through a membrane-embedded rotary motor complex, which subsequently recruits cellular effectors to the activation site to degrade the phage genome (Hu *et al*., 2025; Mariano *et al*., 2025). Zorya II encodes a single effector protein ZorE, which has been suggested to function as a nickase (Mariano *et al*., 2025). This raises the possibility that the synergistic effect of the two systems may be achieved at the level of intracellular effectors, with ZorE cleaving the phage DNA, unwound by DruE.

In summary, we provide functional and structural insights into a minimal two-protein module Druantia anti-phage defense system and present a model describing how DruE coordinates with DruH to achieve a direct antiviral response.

## Methods

### Bioinformatical analysis

Pre-annotated bacterial defense systems from RefSeq release 209 were downloaded from PADLOC-DB v2.0.0 (Payne *et al*., 2022) and used to estimate the abundance of Druantia systems. Genomes containing Druantia hits were retained for downstream phylogenetic analysis. Open reading frames encoding DruE and DruH were extracted and clustered by sequence similarity using MMseqs2 (v 16-747c6)(Steinegger and Söding, 2017): mmseqs easy-cluster input.fa output 05 tmp --min-seq-id 0.50-c 0.8 --cov-mode 1. Cluster representative sequences were used as input for multiple sequence alignment with MAFFT (v7.525) (Katoh and Standley, 2013): mafft --maxiterate 100 --globalpair input.fa > output.fa. Alignments were trimmed using trimAl (v1.5.rev0) (with flag-gt 0.25) (Capella-Gutiérrez, Silla-Martínez and Gabaldón, 2009) and phylogenetic trees were inferred using IQ-TREE (v3.0.1)(Wong *et al*., 2025): iqtree-s input-bb 2000-alrt 1000-m MFP+MERGE-pre out. Annotation of the putative dimerization domain was performed using HMMER v3.4 (hmmbuild and hmmsearch) (Eddy, 2011). Briefly, representative sequences corresponding to the dimerization domain were selected and aligned using Clustal Omega (v1.2.4)(Sievers and Higgins, 2018). The resulting multiple sequence alignment was used to build a hidden Markov model (HMM) profile, which was subsequently used to identify homologous regions across all PADLOC DruE proteins: hmmsearch --domtblout out --incE 1e-2 --incdomE 1e-2 input.hmm database.fa. Phylogenetic trees were visualized using pyCirclize (v1.2.0) and Biopython (v1.85). All analyses were conducted in Python v3.11.9 using pandas (v2.1.1), and Matplotlib (v3.7.1).

### Cloning of *Ec*DruIII defense system and mutagenesis

The full-length genes of *E. coli DruH* and *DruE* code for 1119 and 2104 residues, respectively. The *Ec*DruIII operon with its native promoter region was PCR amplified from the *E. coli* strain NCTC12923 genome (purchased from the National Collection of Type Cultures, NCTC) and subcloned into a modified pET vector using In-Fusion cloning strategy (In-Fusion® Snap Assembly Master Mix; TaKaRa Cat. # 638947). To facilitate purification and cleavage, a TEV protease recognition site and a 6*His tag was incorporated before *DruH* gene, and a 3C protease recognition site and a twin-Strep-tag II (TSII) were incorporated after *DruE* gene, resulting in the construct: *pET11a-T7-promoter-Dru-native-promoter_6×His-TEV-DruH-DruE-3C-TSII*. To improve the purity of DruH protein, we PCR amplify *DruH* gene alone and subcloned into the pET vector resulting in the construct: *pET11a-T7-DruH-3C-TSII*. For generating mutations (point mutations, deletions) plasmids were constructed based on standard cloning techniques (In-fusion snap assembly) or purchased from Genscript. All plasmids were verified by either Sanger or Nanopore sequencing.

### Phage infectivity assays

The host strain *E. coli* K-12 ΔRM (hereafter K-12ΔRM, a derivative of *E. coli* K-12 engineered to remove multiple restriction modification systems, which was used isolate the BASEL phage collection (Maffei *et al*., 2021)), possessing either pControl (pET11a with *Ec*DrIII native promotor) or p*Ec*DrIII (or mutants thereof) were grown overnight in LB + Ampicillin (Amp^+^; 100 µg/mL). Efficiency of plaquing (EOP) assays were performed using bacterial lawns of the host strain (OD: 0.3) in (1:50 dilution) 0.5% LB agar + 10 mM MgSO_4_ + 2 mM CaCl_2_ overlaid onto 1.5% LB agar + Amp^+^. Ten-fold dilution series of phages were spotted onto the overlays, air-dried, then the plates were incubated overnight at 30°C. Efficiency of plating (EOP) was measured as the total number of plaque-forming units (PFUs) on the experimental strain carrying control plasmids, divided by the number of PFUs on the strain carrying the wildtype (WT) or mutant plasmids. Plaques were counted at the dilution where individual plaques were countable. If phage infection resulted in a lysis zone without distinguishable plaques (i.e., no discrete plaques), the last dilution showing lysis was counted as having 10 plaques.

### Population growth assay

Overnight cultures were grown in LB with respective antibiotics. Subsequently, cultures were diluted in fresh LB-Lennox medium supplemented with antibiotics, MgSO_4_ and CaCl_2_ to an OD600 of 0.05. Cells were then split in a 96-well-plate (transparent walls, clear bottom, Greiner Bio-One, Frickenhausen, Germany) and exposed to phages at indicated MOIs. The assay was performed using a H1 plate reader (BioTek, Winooski, VT, USA). The optical density was measured every 5 min for at least 18h under constant aeration at 37°C.

### Phage DNA purification

Phage primary stocks were propagated using the double-layer agar method with 0.6% soft agar and 1% hard agar. Agar overlays were supplemented with 5 mM MgCl₂ and poured using E. coli K-12ΔRM cultures grown at 37°C to an OD₆₀₀ of 0.6. Plates were incubated overnight at 37°C until plaques formed. Phage lysates were recovered by adding 10 mL of SM buffer pH 7.5 (composition: 100 mM NaCl, 10 mM MgSO_4_, 50 mM Tris-HCl) to each plate and incubating overnight at 4°C with gentle shaking. The resulting supernatant was collected and clarified by addition of chloroform (final concentration 1% v:v), followed by centrifugation at 4,000 × g for 15 min at 4°C. The clarified lysate was treated with DNase I at 37°C, and the enzyme was subsequently heat-inactivated at 75°C for 5 min. Phage genomic DNA was then purified using the Norgen Phage DNA Isolation Kit (Norgen Biotek, Thorold, ON, Canada) according to the manufacturer’s instructions.

### Protein expression and purification

#### DruE

The plasmid containing wildtype *Ec*DruIII operon: *pET11a-T7-promoter-Dru-native-promoter_6×His-TEV-DruH-DruE-3C-TSII*. were transfected into *E. coli* BL21 star (DE3) pRARE competent cells and the proteins were expressed in LB medium. When the culture OD_600_ reached to 0.6-0.8, the temperature was decreased from 37°C to 18°C, then grown until the OD_600_ reached approximately 0.8-1.0, and 0.5 mM isopropyl β-D-1-thiogalactopyranoside (IPTG) was added for overnight protein induction. The culture was harvested, and the cell pellet was resuspended in buffer A containing 20 mM HEPES-NaOH pH 7.5, 300 mM NaCl, 10% Glycerol, 1mM TCEP supplemented with EDTA-free protease inhibitor (Thermo Fisher Scientific) and lysozyme from chicken white egg (Sigma) to a final concentration of 50 μg/mL. The mixture was disrupted by high-pressure homogenizer and spun at 18,000 g for 1 h. The supernatant was loaded onto a gravity flow column containing 2 mL (resin volume) of Strep-Tactin® 4flow® high-capacity resin (IBA), pre-equilibrated with wash buffer containing 20 mM HEPES-NaOH pH 7.5, 300 mM NaCl, 10% glycerol, 1mM TCEP. The resins were washed five times with 2-3 resin volumes of the wash buffer and elution was carried out five times with 0.5 resin volume (1 mL) of elution buffer containing 20 mM HEPES-NaOH pH 7.5, 300 mM NaCl, 10% glycerol, 1mM TCEP and 10 mM desthiobiotin. The recombinant protein was then concentrated and loaded onto a pre-equilibrated (20 mM HEPES-NaOH pH 7.5, 150 mM NaCl, 1mM TCEP) Superose 6 Increase 10/300 GL size exclusion chromatography column. Fractions from the elution peak corresponding to the molecular weight of DruE protein were pooled, and the protein was concentrated for cryo-EM grid preparation and functional experiments. The procedures of expression and purification of DruE mutants were similar as the DruE wildtype.

#### DruH

The predicted DruH gene encodes a protein of 1,119 amino acid residues. We initially purified DruH from the functional plasmid *pET11a-T7-promoter-Dru-native-promoter_6×His-TEV-DruH-DruE-3C-TSII* using a His-tag affinity purification approach. However, the purity was suboptimal due to nonspecific binding of *E. coli* proteins to the Ni-NTA resin. We therefore switch to the plasmids *pET11a-T7-DruH-3C-TSII*, and purify DruH using Twin-Strep tag. DruH purification steps were similar as DruE. Briefly, the suspension buffer contained 300 mM NaCl, 20 mM HEPES-NaOH pH 7.5, 1 mM TECP and 10% glycerol; the wash buffer was the same as the suspension buffer and the elution buffer contained 150 mM NaCl, 20 mM HEPES-NaOH pH 7.5, 10% glycerol, 1 mM TCEP and 10 mM desthiobiotin; and the size exclusion chromatographic buffer contained 150 mM NaCl, 20 mM HEPES-NaOH pH 7.5 and 1 mM TCEP. Fractions from the elution peak corresponding to the molecular weight of DruH protein were pooled, and the protein was concentrated for cryo-EM grid preparation and functional experiments. The procedures of expression and purification of DruH mutants were similar as the DruH wildtype.

### Mass spectrometry trypsin in-gel digestion sample preparation, data acquisition and analysis

Proteins were separated using precast 4–20% Tris-Glycine SDS-PAGE gels (1.0 mm thick; Life Technologies, Carlsbad, CA). The gel was stained with SimplyBlue SafeStain (Life Technologies, Carlsbad, CA). Protein bands of interest were excised from the polyacrylamide gel, destained, and dehydrated prior to reduction, alkylation, and in-gel tryptic digestion. The gel pieces were reduced with dithiothreitol (43815, Sigma) and alkylated with iodoacetamide (I1149, Sigma). Subsequentely, the gel pieces were dried and rehydrated in 50 mM ammonium bicarbonate (09830, Sigma), 12 ng/μL trypsin (mass spectrometry grade; V5111, Promega), and 0.01% ProteaseMAXTM surfactant (trypsin enhancer; V2071, Promega). After a 10 min incubation at room temperature, the rehydrated gel pieces were overlaid with 30 μL of 0.01% ProteaseMAX in 50 mM ammonium bicarbonate and incubated at 37°C for 3 hours with shaking at 300 rpm on a Thermomixer (Eppendorf). The digestion mixture was transferred to a fresh tube, acidified with formic acid to a final concentration of 1%, and centrifuged at 14,000 x g for 10 min to remove particulate material. The supernatant was stored at-20°C until LC-MS/MS analysis.

The tryptic peptides were separated on a Hypersil GOLDTM C18 column (100 mm × 1 mm, 3 μm particle size, 175 Å pore size; 25003-101030, Thermo Fisher Scientific) using an Ultimate 3000 LC system (Dionex, Thermo Fisher Scientific). The mobile phases composition was as follows: A: 0.1% (v/v) formic acid in water with 2% acetonitrile and B: 98% acetonitrile, 2% water, and 0.1% (v/v) formic acid. The gradient program was: 0 min, 5% B; 5 min, 11% B; 30 min, 25% B; 45 min, 50% B. The flow rate was 0.15 ml/min, and the column oven temperature was maintained at 45°C. The ions were infused into a micrOTOF-QII mass spectrometer (Bruker Daltonics GmbH & Co.) equipped with an electrospray ionization (ESI) source operated in positive mode. The ESI settings were: capillary voltage 4500 V, end plate offset-500 V, nebulizer gas (nitrogen) pressure 1.4 bar, drying gas flow 9 L/min, and drying gas temperature 190 °C. A precursor m/z scan range of 75–2200 m/z was used, followed by data dependent MS/MS acquisition of the top five most abundant precursor ions in each full MS1 scan.

Data were analyzed using Bruker Compass DataAnalysis version 6.0 (Bruker Daltonics GmbH & Co.). Mass calibration was performed using sodium formate cluster ions as an internal calibrant. Peptide identification was carried out using AutoMS(n) with a signal intensity threshold of 1000 and spectra were deconvoluted using the “Peptides/Small Molecules” parameter preset. The detected peptides were submitted to an automated Mascot search for identification against the Swissprot database and the druantia E or H construct sequences (Perkins *et al*., 1999). In-silico digestion was perfomed using “Trypsin/P” allowing up to two missed cleavages. Carbamidomethylation of cysteine residues was set as a fixed modification and protein N-terminal acetylation, oxidation of methionine and deamidation of asparagine and glutamine were included as potential variable modifications. The search mass tolerance was set to MS: 10 ppm and MS/MS: 0.05 Da.

### Size exclusion chromatography coupled with multi-angle static laser light scattering (SEC-MALS)

SEC-MALS experiments were performed on a Dionex (Thermo ScientificTM) HPLC system connected in-line with a UV detector (Thermo ScientificTM DionexTM Ultimate 3000, MWD-3000), a DAWN HELEOS 8+ multiangle laser light scattering detector and an Optilab T-rEX (Wyatt Technology Corporation) refractive index detector. SEC was performed on a SuperdexTM 200 Increase 10/300 GL column (GE Healthcare) at room temperature in 20 mM Hepes buffer pH 7.5, 150 mM NaCl, 1.0 mM TCEP. 100 μl of DruE were injected at a series of concentrations between 0.06 mg/ml and 4 mg/ ml and at a flow rate of 0.5 ml/min. 100 μl of the DruE PD mutant were injected at a single concentration, 2 mg/ml and at 0.5 ml/ min. The ASTRA (version 8.0.2.5 64-bit) software (Wyatt Technology Corporation) was used to collect the data from the ultraviolet, refractive index and light scattering detectors. The weight average molecular masses were determined across the elution profiles from static LS measurements using ASTRA software and a Zimm model which relates the amount of scattered light to the weight average molecular weight, the concentration and the square of the refractive index increment (dn/dc) of the sample.

### Determination of DruE dimerization equilibrium constant from SEC-MALS

The concentration-dependent changes in the weight-average molecular mass determined at the apex of the DruE eluting peak were used to calculate the dimerization constant. The weight-average molecular masses from three independent concentration series were plotted as a function of DruE concentration and fitted to a monomer-dimer equilibrium using a nonlinear least-squares algorithm.

### Mass photometry (MP)

Mass photometry experiments were performed on a Refeyn OneMP (Refeyn Ltd) mass photometer at room temperature. Measurements were run in triplicates diluting 4 μl of protein samples at 40 nM in 12 μl in 20 mM Hepes buffer pH 7.5, 150 mM NaCl, 1.0 mM TCEP loaded on a gasket (Grace Bio-Labs reusable CultureWellTM gaskets, Merck (#GBL103250)) mounted on a glass microscope coverslip (Marienfeld (#0107222)). The molecular weights were obtained by contrast comparison with known molecular weight mass standard calibrants (NativeMark Unstained Protein Standard, ThemoFisher Scientific, (#LC0725)) measured on the same buffer and on the same day. Movies were recorded using AcquireMP (Refeyn Ltd, version 2025 R1.2), with a medium field of view and exposure time of 0.95 ms for the acquisition camera settings. A total of 6000 frames were recorded over a duration of one minute. Movies were processed and analysed with the DiscoverMP software (Refeyn Ltd, version 2025 R1) provided by the instrument manufacturer.

### Malachite green ATPase assay

ATP hydrolysis was measured using a time-course ATPase assay and Malachite Green Phosphate Detection Kit (“Cell Signaling Technology”). Reactions were carried out in reaction buffer (20 mM Tris-HCl pH=7.5, 50 mM NaCl, 1 mM TCEP and 1mM MgCl_2_). Where specified, reactions were supplemented with 500 µM ATP. Samples were pre-incubated for 2 min at 37°C and DruE, its mutant or DruH was added to initiate the reaction. Reactions were performed in a total volume of 25 µL containing 100 nM of protein with or without 1 µM of ssDNA (5‘-GGTACCCTTGAGGTGCTGTTTCAGGGACCAGGAGGTAGTGTACTGAAGTTCAAACA AAT-3’). ATP hydrolysis was monitored over time by removing 5 µl aliquots at 5, 10, 20, and 30 minutes and mixing with 5 µl of stop solution (10 mM EDTA). Released inorganic phosphate (P_i_) was quantified using Malachite Green Phosphate Detection Kit (“Cell Signaling Technology”) according to the manufacturer instructions. 96-well flat bottom plate was scanned on a plate reader instrument (Spark Multimode Microplate Reader, Tecan) at 630 nm wavelength. The data were visualized using the Prism software.

### DruE helicase assays

5’FAM-labeled ssDNA and unlabeled complementary DNA were mixed at a 1:2 molar ratio, denatured at 95 °C for 5 min, and gradually cooled to ambient temperature. The DNA annealing buffer contained 10 mM Tris-HCl (pH 7.5) and 50 mM NaCl. **Table 2** lists the sequences of the dsDNA substrates: 3’overhang, 5’overhang, blunt-ended, and forked DNA. For the helicase assay, 100 nM 5’FAM-labeled dsDNA substrates were incubated with DruE or DruE mutants at the indicated concentrations (50 nM, 500 nM, 1 µM) in the presence of 100 µM ATP or AMP-PNP in helicase buffer. A 5× helicase buffer was prepared containing 100 mM Tris-HCl (pH 7.5), 250 mM KCl, 5 mM MgCl₂, 50% (v/v) glycerol, 5 mM DTT, and 500 µg ml⁻¹ BSA. Additionally, 5 µM unlabeled ssDNA (corresponding to the 5’FAM-labeled strand) was included to prevent reannealing. Reactions were incubated at 30 °C for 15 min, after which 1× stop buffer was added. A 5× stop solution was prepared containing 2 mg/ml proteinase K (New England Biolabs), 90 mM EDTA, 1.8% SDS, and 50% glycerol. Samples were loaded onto a 10% TBE polyacrylamide gel and electrophoresed in 1× TBE buffer at 200 V for 30-45 min at 4 °C. The gels were visualized using Odyssey^®^ XF Imaging System at 600 nm.

### DruH electromobility shift assays (EMSA)

Frozen aliquots of DruH in size exclusion buffer (150 mM NaCl, 20 mM HEPES-NaOH pH 7.5 and 1 mM TCEP) were thawed and centrifuged to remove aggregates and diluted to 2x stocks in the same gel filtration buffer. Unless specified otherwise, the final conditions for each reaction consisted of 150 mM NaCl, 20 mM HEPES-NaOH pH 7.5, 1 mM TCEP, 10% glycerol, and 0.5 mM ZnCl2. For each experiment 100 nM of 5’FAM labelled ssDNA substrate was used with the sequence: GGTACCCTTGAGGTGCTGTTTCAGGGACCAGGAGGTAGTGTACTGAAGTTCAAACAAAT from 5’ to 3’. In the case of dsDNA, an unlabeled complementary oligo was annealed to the same substrate. Other ssDNA substrates used for EMSA are listed in **Table S2**. In most cases DruH was added last with a 2.5 µM final concentration (or dilutions thereof in the case of concentration gradients). The reactions were incubated at room temperature for 30 min and loaded on a 1.5% w/v agarose gel made with 20 mM sodium phosphate (pH 7.4). Gels were run for 35 min at 90V at 4°C in 20 mM sodium phosphate (pH 7.4) running buffer and visualized using an Odyssey XF Imaging System at 600 nm.

There were some variations between different EMSA experiments, and the precise details are as follows. For the initial ssDNA vs dsDNA experiment DruH concentrations shown are 0.064, 0.16, 0.4, 1.0, and 2.5 µM. When comparing the various ions, the final concentration of each ion was 50 and 500 µM, and 500 µM for EDTA. The EMSAs comparing NTPs, each consisted of a final concentration of 1 mM NTP, whereas the ATP concentration gradient included 0.0625, 0.25, 1.0, and 4.0 mM. For all of the NTP containing EMSAs DNA was added last, allowing protein and NTP to incubate for 10 min, followed by 20 min incubation with DNA. For denaturing gel, 1.5 ul of each reaction was also run on a denaturing polyacrylamide gel (Novex TBE-Urea, 15%), and again imaged with an Odyssey XF Imaging System at 600 nm.

### Cryo-EM grid preparation, data collection, model building, and refinement

#### DruE apo

Grid preparation and vitrification were performed similarly as previously described (Hu *et al*., 2025). Freshly purified DruE sample was concentrated to 0.8 mg/mL and 2.7 μL protein was applied onto glow-discharged (30 s, 5 mA) grids (UltrAuFoil R0.6/1 300 mesh or UltrAuFoil R1/1 300 mesh) and plunge-frozen into liquid ethane using a Vitrobot Mark IV (Thermo Fisher Scientific), with the settings: 100% humidity, 4°C, blotting force 15-20, 5-7 s blot time and 5 s wait time. Movies were collected using the semi-automated acquisition program EPU (Thermo Fisher Scientific) on a Titan Krios G2 microscope operated at 300 keV equipped with a SelectrisX energy filter and Falcon 4i direct electron detector (Thermo Fisher Scientific). Images were recorded in electron counting mode, at 165,000x magnification with a calibrated pixel size of 0.725 Å and a defocus range of-0.5 to-2.0 μm. The number of micrographs and total exposure values for the different datasets are summarized in Table S1.

#### DruE in complex with forked DNA and ATP

For the DruE complex with forked DNA and ATP, we collected two datasets. For the first dataset, cryo-EM grids were prepared by mixing 4 mg/mL DruE (10 µM, 8 µL), 10 µM DNA substrate (8 µL), and 1 mM ATP. The mixture was incubated at ambient temperature for 3 minutes, after which 10 µM DruH (8 µL) was added, followed by a fourfold dilution. In this dataset, we were only able to resolve cryo-EM maps of the DruE–DNA substrate complex and DruH alone, separately. No complex between DruE and DruH was observed. For the second dataset, cryo-EM grids were prepared by mixing 1.7 mg/mL DruE (10 µM, 8 µL), 10 µM DNA substrate (7 µL), and 1 mM ATP, followed by incubation on ice for 30 minutes. 3 µL of sample were applied onto glow-discharged grids (30 s, 5 mA; UltrAuFoil R0.6/1 300 mesh or UltrAuFoil R1/1 300 mesh) and plunge-frozen in liquid ethane using a Vitrobot Mark IV (Thermo Fisher Scientific) with the following settings: 100% humidity, 4 °C, blotting force 15–20, blot time 5–7 s, and wait time 5 s.

Data collection strategies for the complex were similar to those used for the DruE wildtype. In the second dataset, we were able to resolve cryo-EM maps of the DruE–DNA substrate complex; however, the resolution was lower compared to the first dataset. Therefore, we continued processing only the first dataset to obtain a high-resolution map of DruE with forked dsDNA.

#### DruH

Purified DruH was applied onto glow-discharged grids (30 s, 5 mA; UltrAuFoil R0.6/1 300 mesh or UltrAuFoil R1/1 300 mesh) and plunge-frozen into liquid ethane using a Vitrobot Mark IV (Thermo Fisher Scientific) with the following settings: 100% humidity, 4 °C, blotting force 20, blot time 5–7 s, and wait time 10 s. Data collection strategies for the complex were similar to those used for the DruE wildtype. For the DruH + ssDNA sample, DruH (final concentration 1 mg/mL) was mixed with 10 µM ssDNA. Samples were incubated at room temperature for 10 minutes prior to grid preparation. Samples (3 µL) were applied to UltrAuFoil R1/1 300 mesh Au grids (glow-discharged for 30 s at 5 mA) and plunge-frozen as described above, with the following settings: blotting force 20 and blot time 5–7 s. For DruH in complex with ATP, 10 µL of DruH (final concentration 1.5 mg/mL) was mixed with 10 µL of 10 µM forked dsDNA substrate and 1 mM ATP and incubated at ambient temperature for 10 minutes prior to grid preparation. Data collection strategies for the complex were similar to those described above. In this dataset, we were only able to resolve DruH in complex with ATP.

### Cryo-EM data processing

Cryo-EM data processing steps were performed similarly as previously described(Hu *et al*., 2025). All datasets were processed using cryoSPARC (Punjani *et al*., 2017) v4.7, unless otherwise stated. We started by using Patch motion correction to estimate and correct for full-frame motion and sample deformation (local motion). Patch Contrast Function (CTF) estimation was used to fit local CTF to micrographs. Micrographs were curated to remove low-quality data (the relative ice thickness value greater than 1.1 and the CTF fit value worse than 3.5 Å). We performed particle picking by template picking on denoised micrographs. Particles were extracted with a box size of 500^2^ pixels for DruE datasets, and 400^2^ pixels for the DruH datasets. One round of 2D classification was performed followed by *ab initio* reconstruction. Several rounds of heterogeneous refinement were used to exclude broken particles. Non-uniform refinement was applied with a dynamic mask to obtain a high-resolution map. Local refinement was additionally performed with a soft mask to achieve a higher-resolution map of some flexible regions. For all datasets, the number of movies, the number of particles used for the final refinement, map resolution, and other values during data processing are summarized in the **Table S1**.

### Model building and validation

We used AlphaFold3 (Abramson *et al*., 2024) to predict the initial models of DruE and DruH apo structures. The predicted models were manually fit into the cryo-EM density by using UCSF ChimeraX (Pettersen *et al*., 2021). The models were then refined in Coot (Emsley and Cowtan, 2004) or using StarMap (Lugmayr *et al*., 2023). DNA substrates, including the ideal B-formed dsDNA, were generated using Coot. The models were then refined against the map using PHENIX real space refinement (Liebschner *et al*., 2019). The DruE PLD1 and PLD2 domains in the reconstructions of DruE dimer and in complex with the forked dsDNA are flexible and were manually adjusted to best fit the density. For structural comparison using the Z-score, the calculation was performed similarly to that in a previous study (Hu *et al*., 2023).

### Mass spectrometry sample preparation

Mass spectrometry sample preparation steps were performed similarly as previously described(Hu *et al*., 2025). Overnight cultures of *E. coli* K-12 ΔRM transformed with p*Ec*DruIII (or pControl) were used to inoculate (at a 1:100 dilution) 3 ml LB medium with antibiotics, then grown to an OD600 of approximately 0.4. The cells were pelleted and resuspended in 750 µl HBS buffer (20 mM HEPES pH 7.5, 150 mM NaCl, 1 mM TCEP). Next, 250 µl of 6% (w/v) SDS was added to reach a final concentration of 1.5%, and the sample was heated at 99 °C for 10 min. The mixture was then sonicated (Misonix Ultrasonic Liquid Processor with microtip probe) to fragment DNA and RNA using the following settings: amplitude 10; 5 s sonication followed by 5 s pause; 5 cycles.

For MS analysis, we performed protein aggregate capture digestion(Batth *et al*., 2019). To this end, 250 µl of bacterial lysate was taken from the total sample, and 750 µl of acetonitrile was added, along with 50 µl of magnetic microspheres prewashed with PBS. The mixture was allowed to settle for 10 min before collecting the magnetic microspheres on a magnetic plate. The beads were washed once with 1 ml acetonitrile and once with 1 ml 70% ethanol. After removing all ethanol, the beads were stored at −20 °C until further processing.

Frozen beads were thawed on ice, supplemented with 100 µL ice-cold 50 mM TRIS pH 8.5 buffer supplemented with 2.5 ng/µL trypsin, and gently mixed (on ice) every 5 min for 30 min. Digestion was performed for 2 h using a ThermoMixer shaking at 1,250 rpm at 37 °C. Peptides were separated from magnetic microspheres using 0.45 µm filter spin columns, and peptides were reduced and alkylated by adding TCEP and chloroacetamide to 5 mM for 30 min prior to peptide clean-up via low-pH C18 StageTip procedure. C18 StageTips were prepared in-house, by layering four plugs of C18 material (Sigma-Aldrich, Empore SPE Disks, C18, 47 mm) per StageTip. Activation of StageTips was performed with 100 μL 100% methanol, followed by equilibration using 100 μL 80% acetonitrile (ACN) in 0.1% formic acid, and two washes with 100 μL 0.1% formic acid. Samples were acidified to pH <3 by addition of trifluoroacetic acid to a concentration of 1%, after which they were loaded on StageTips. Subsequently, StageTips were washed twice using 100 μL 0.1% formic acid, after which peptides were eluted using 80 µL 30% ACN in 0.1% formic acid. All fractions were dried to completion using a SpeedVac at 60 °C. Dried peptides were dissolved in 50 μL 0.1% formic acid (FA) and stored at −20 °C until analysis using mass spectrometry (MS).

Approximately 1 µg of peptide was analyzed per injection. All samples were analyzed on a Vanquish™ Neo UHPLC system (Thermo Fisher Scientific) coupled to an Orbitrap™ Astral™ mass spectrometer (Thermo Fisher Scientific). Samples were analyzed on 20 cm long analytical columns, with an internal diameter of 50 μm, and packed in-house using ReproSil-Pur 120 C18-AQ 1.9 µm beads (Dr. Maisch). The analytical column was heated to 40 °C, and elution of peptides from the column was achieved by application of gradients with stationary phase Buffer A (0.1% FA) and increasing amounts of mobile phase Buffer B (80% ACN in 0.1% FA). The primary analytical gradient ranged from 10 %B to 38 %B over 42.5 min, followed by a further increase to 48 %B over 5 min to elute any remaining peptides, and by a washing block of 7.5 min. Ionization was achieved using a NanoSpray Flex NG ion source (Thermo Fisher Scientific), with spray voltage set at 2 kV, ion transfer tube temperature to 275 °C, and RF funnel level to 50%. All full precursor (MS1) scans were acquired using the Orbitrap™ mass analyzer, while all tandem fragment (MS2) scans acquired in parallel using the Astral™ mass analyzer. Full scan range was set to 300-1,000 m/z, MS1 resolution to 180,000, MS1 AGC target to “500” (5,000,000 charges), and MS1 maximum injection time to 250 ms. Precursors were analyzed in data-dependent acquisition (DDA) mode, with charges 2-6 selected for fragmentation using an isolation width of 1.3 m/z and fragmented using higher-energy collision disassociation (HCD) with normalized collision energy of 25. Monoisotopic Precursor Selection (MIPS) was enabled in “Peptide” mode. Repeated sequencing of precursors was minimized by setting expected peak width to 15 s, and dynamic exclusion duration to 15 s, with an exclusion mass tolerance of 10 ppm and exclusion of isotopes. MS2 scans were acquired using the Astral mass analyzer. MS2 fragment scan range was set to 100-1,500 m/z, MS2 AGC target to “100” (10,000 charges), MS2 intensity threshold to 50,000 charges per second, and MS2 maximum injection time to 10 ms; thus requiring a minimum of 500 charges for attempted isolation and identification of each precursor. Duty cycle was fixed at 0.5 s.

### MS data analysis

All RAW files were analyzed using MaxQuant software (version 2.5.2.0. Default MaxQuant settings were used, with exceptions outlined below. For generation of the in silico spectral library, the two full-length Druantia protein sequences and the mNeonGreen sequence were entered into a FASTA database, along with all (23,272) Swiss-Prot-reviewed Escherichia coli sequences (taxonomy identifier 562) downloaded from UniProt on the 18th of December, 2025. The tryptic peptide resulting from the Strep-tag (GGGSGGGSGGSAWSHPQFEK) was included as a separate entry to prevent quantification artefacts. For the main data search using MaxQuant, digestion was performed using “Trypsin/P” with up to 4 missed cleavages, with a minimum peptide length of 6 and a maximum peptide mass of 5,000 Da. No variable modifications were considered for the first MS/MS search, which is only used for precursor mass recalibration. For the MS/MS main search a maximum allowance of 3 variable modifications per peptide was set, including oxidation of methionine (default), deamidation of asparagine, and peptide N-terminal glutamine to pyroglutamate. N-terminal protein acetylation was disabled as a default variable modification, as this does not occur in prokaryotes. Modified peptides were stringently filtered by setting a minimum score of 20, and a minimum delta score of 40. First search mass tolerance was set to 10 ppm, and maximum charge state of considered precursors to 6. Label-free quantification (LFQ) was enabled, “Fast LFQ” was disabled. iBAQ was enabled. Matching between runs was enabled with a match time window of 0.5 min and an alignment time window of 10 min. Data was filtered by posterior error probability to achieve a false discovery rate of <1% (default), at the peptide-spectrum match, protein assignment, and site-decoy levels.

### Mass spectrometry data statistics

All statistical data handling was performed using the Perseus software (Tyanova *et al*., 2016), including data filtering, log2-transformation, imputation of missing values (down shift 1.8 and width 0.15), and two-tailed two-sample Student’s t-testing with permutation-based false discovery rate control. In order to determine relative concentration of all proteins in the samples, LFQ-normalized intensity values for each protein were adjusted by molecular weight. To approximate absolute copy numbers, we extracted known protein copy numbers based on the “LB” condition as reported by Schmidt et al. (Schmidt *et al*., 2016), log2-transformed them, and aligned them to the molecular-weight adjusted LFQ intensity values from our own data, resulting in 1,702 out of 2,082 quantified protein-groups receiving a known copy number value (R² = 0.5906). Next, we subtracted the overall median from all log2 values and determined the absolute delta between the values of each pair. Out of all pairs, 348 had a log2 delta of <0.5, which we considered as a “proteomic ruler”. Linear regression was performed on the remaining pairs (R² = 0.9866) to determine a conversion factor between MW-adjusted LFQ intensity and absolute copy number.

### Single-cell time-lapse microscopy growth assay

Overnight cultures of *E. coli* strains carrying *Ec*DruIII plasmids were grown with shaking at 180 rpm in LB Lennox medium containing 10 mM MgSO₄ and 2 mM CaCl₂, supplemented with 100 µg ml⁻¹ ampicillin at 30 °C. The following day, a 1:100 subculture was inoculated and grown at 30 °C until an OD₆₀₀ of 0.3–0.5 was reached. Cultures were then diluted to an OD₆₀₀ of 0.2. Cells were exposed to phages (Bas08 or Bas08 *parS* inserted after MCP) at indicated MOI for 5 min (or left untreated) under shaking conditions. For the endpoint measurement, cells were incubated until the indicated timepoint. Subsequently, 1 µl of the culture was spotted onto a 1.2% (w/v) agarose pad (prepared in LB:MQ at a 1:5 ratio). Microscopy slides were mounted in an incubation chamber preheated to 30 °C. Image acquisition was performed using a Nikon Eclipse Ti2 inverted microscope equipped with a CFI Plan Apochromat DM ×60 Lambda oil Ph3/1.40 objective. Phase-contrast images were captured every 2 min for the indicated duration. Data analysis was performed with customized python scripts.

### Bacterial genome extraction and Nanopore sequencing

Overnight cultures of *E. coli* K-12 ΔRM transformed with p*Ec*DruIII (or pControl; Vector-mNg) were used to inoculate 3 mL LB medium containing antibiotics at a 1:100 dilution and grown to an OD600 of approximately 0.4. Genomic DNA was extracted using the GenElute Bacterial Genomic DNA Kit (Sigma-Aldrich; NA2100-1KT) according to the manufacturer’s instructions. The genomic DNA, without additional fragmentation, was end-repaired with the addition of dA overhangs, followed by T/A ligation of barcodes. Samples were sequenced on an R10.4.1 nanopore (Kit 14 chemistry). Dorado v1.0.0 was used for base calling with the super-accuracy v5.2.0 model, including the 6mA, 4mC_5mC, and 5mC_5hmC modification models. 5mC modification was evaluated using two different models, resulting in two independent measurements that were slightly different, which is expected. Modification analysis was performed using modkit v0.6.1. The results were first evaluated using the “modkit summary” function to determine differences in global methylation, specifically whether the case sample contained considerably more methylated sites compared to the control.

### Construction of the phage Bas08-parS

We introduced a *parS* site downstream of the major capsid protein gene of *Escherichia* virus DanielBernoulli (Bas08) (Maffei *et al*., 2021) by homologous recombination with a synthetic template followed by CRISPR-Cas9 selection against the parental wildtype using a setup similar to the procedure described elsewhere (Esvelt *et al*., 2013). To generate the template for homologous recombination, we used PCR to insert the *parS* sequence (TCGCCATTCAAATTTCACTATTAACTGACTGTTTTTAAA GTAAATTACTCTAAAATTTCAAGGTGAAATCGCCACGATTTCAC) between plasmid-encoded homology arms of ca. 400 bp length by PCR (Liu and Naismith, 2008), generating plasmid pAH210_Bas08-MCP_parS. To prevent CRISPR activity against the recombinant phage, silent point mutations were introduced into the PAM and protospacer inside the *mcp* gene while preserving the local amino acid sequence. Subsequently, we transformed an *E. coli* K-12 with this plasmid and then infected it with phage Bas08 to enable homologous recombination, generating a lysate that would contain both wildtype and recombinant phages. We then used CRISPR-Cas9 with a crRNA targeting GAATCAGGATCATATTAGCG (5’-3’) in the wildtype *mcp* gene to select for the desired recombinant clones. CRISPR-Cas9 selection was performed using a two-plasmid setup of DS-SPcas (expressing *cas9*) and pAH212_crispr-Bas08-MCP (expressing the crRNA). Finally, plaques obtained after CRISPR-Cas9 selection were screened for successful insertion of the *parS* sequence and absence of undesired mutations site by Sanger sequencing of PCR products spanning the whole targeted region.

### Figure preparation

Structural figures were prepared using ChimeraX version 1.10 (Pettersen *et al*., 2021), GraphPad Prisim10 and Adobe Illustrator (*Adobe Illustrator*, 2021).

## Data and Code Availability

Atomic coordinates for DruE apo monomer, DruE apo dimer, DruE dimer in complex with forked dsDNA were deposited in the Protein Data Bank (PDB) under accession codes 28YG, 28YS and 28ZH, respectively. The corresponding electrostatic potential maps were deposited in the Electron Microscopy Data Bank (EMDB) under accession codes EMD-56965, EMD-56979 and EMD-57002, respectively. The local refinement map of PD subdomain in DruE apo dimer were deposited in the EMDB under accession codes EMD-. Atomic coordinates for DruH apo, DruH in complex with ssDNA, DruH in complex with ATP were deposited in the PDB under accession codes 28XY, 28YA and 28YB, respectively. The corresponding electrostatic potential maps was deposited in the EMDB under accession codes EMD-56950, EMD-56956 and EMD-56957. The mass spectrometry proteomics data have been deposited to the ProteomeXchange Consortium via the PRIDE (Perez-Riverol *et al*., 2025) partner repository with the dataset identifier PXD075049. Project accession: PXD075049. Token: IxTgLVqBQkhD

## Acknowledgements

The Novo Nordisk Foundation Center for Protein Research is supported financially by the Novo Nordisk Foundation (NNF14CC0001). N.M.I.T. acknowledges support from an NNF Hallas-Møller Ascending Investigator grant (NNF23OC0081528). N.M.I.T. is also a member of the Integrative Structural Biology Cluster (ISBUC) at the University of Copenhagen. V.K.S and N.N.R. acknowledge support from Novo Nordisk Bioscience Ph.D. program, NNF0069780 and NNF0078229, respectively. We thank the Danish Cryo-EM Facility at the Core Facility for Integrated Bioimaging (CFIB) at the University of Copenhagen for support during data collection.

P.F.P and M.E. acknowledge funding from the Deutsche Forschungsgemeinschaft (DFG) in the framework of the priority program SPP2330 (research grant no. 548567920 to P.F.P (PO 2831/2-1) and to M.E. (ER 778/13-1)).

## Author contribution

H.H. and N.M.I.T. conceived the project. H.H. did molecular biology and mutagenesis, expressed, purified, optimized, and prepared cryo-EM grids, collected the cryo-EM data, and determined all the structures presented in this study. H.H. performed phage infectivity assays. P.F.P. carried out and analyzed liquid culture infection time courses assays, single-cell microscopy experiments with the analysis and together with M.E. interpreted the data. N.R.R. performed all the EMSA experiments. B.L.M. performed mass photometry, SEC-MALS, in gel digestion coupled mass spectrometry assays. V.K.S. performed bioinformatic analyses. A.R-E. and V.K.S. purified phage and phage genomes. I.S. purified *E. coli* genomes and performed ATPase assays. N.H.S. and T.P. assisted during the cryo-EM data collection. H.H. prepared samples for Nanopore sequencing.

I.A.H. performed single cell mass spectrometry and together with J.V.O. analyzed the data. A.H. and D.P. designed and engineered the Bas08_parS phage construct. H.H. prepared figures and wrote the first draft of the manuscript with input from all the authors. All authors contributed to the revision of the manuscript.

## Supplementary Figure Legends

**Figure S1. DruE exits as monomer and dimer and cryo-EM dataset processing results**

(A) Schematic representation of the cloning strategy for *Ec*DruIII.

(B) Plaque assays of phages challenging *E. coli* K-12 ΔRM expressing either a vector control or *Ec*DruIII. Images are representative of three replicates.

(C) Representative SDS-PAGE gel of the co-purifying DruH and DruE proteins. The first lane shows co-purification using a His-tag on DruH, and the second lane shows co-purification using a twin Strep-tag on DruE. Neither affinity-purified DruH nor affinity-purified DruE pulled down the other protein. Images are representative of three replicates.

(D) Mass photometry analysis of purified DruE at 10 nM concentration. The data shown represent combined distributions from three independent measurements.

(E) Cryo-EM image of the DruE sample under cryogenic conditions.

(F) Representative 2D class average of the DruE monomer and dimer.

(G) 3D classes of DruE apo dataset with particle populations.

(H) Cryo-EM density map of the DruE monomer colored by local resolution (Å), estimated in cryoSPARC, and gold-standard (0.143) Fourier shell correlation (GSFSC) curves of the refined complex with particle angles and poses.

(I) Cryo-EM density map of the DruE dimer colored by local resolution (Å), estimated in cryoSPARC and gold-standard (0.143) Fourier shell correlation (GSFSC) curves of the refined EM map with particle angles and poses.

**Figure S2. DruE structure and domains.**

(A) Zoom-in view of the DruE structures, showing that the DruE NTR spatially locks the core folds.

(B) Amino acids and secondary structure of the DruE PD subdomain with part of the Cap domain. Peptides identified by mass spectrometry that cover the DruE PD subdomain are indicated as black lines beneath the residues.

(C) Superimposition of DruE PLD1 and PLD2 with the top hit (PDB ID: 7CLG) from Foldseek.

(D) Phylogeny of DruIIIE sequences, with the presence of PLD1 and/or PLD2 subdomain indicated.

**Figure S3. Structure details of DruE PD subdomains, core domains.**

(A) Gold-standard (0.143) Fourier shell correlation (GSFSC) curves of the refined DruE PD subdomain from the DruE dimer, with particle angles and poses.

(B) Cryo-EM density map and model of the dimerized DruE PD subdomain.

(C) Zoomed-in view of the dimerized PD subdomain interface.

(D) Phylogeny of DruE sequences, with HMM hits for the dimerized PD subdomain indicated across Durantia families.

(E) Structural comparison of the DruE monomer with protomer A in the DruE asymmetric dimer.

(F) Structural comparison of the DruE monomer with protomer B in the DruE asymmetric dimer. In (E) and (F), the PD subdomain, PLD1, and PLD2 are excluded.

(G) Close-up view of the interactions at domain assembly interface III, mediated by Y191 from protomer A and three histidines from protomer B, with the cryo-EM map overlaid.

(H) Size-exclusion chromatography (SEC) of the purified Y191A mutant in comparison with DruE WT, DruE ΔK683-N753 (ΔPD) and DruE DEAH mutant (D252A/E253A/H254A).

**Figure S4. Structure of DruE in complex with forked dsDNA**

(A) Malachite green ATPase assay of DruE, DruE mutant or DruH with or without ssDNA.

(B) Representative SDS-PAGE gel of the purified DruE proteins used for helicase assays. Images are representative of at least three replicates.

(C) Cryo-EM density map of the DruE dimer in complex with forked dsDNA, colored by local resolution (Å) estimated in cryoSPARC, and gold-standard (0.143) Fourier shell correlation (GSFSC) curves of the refined complex with particle angles and poses.

(D) Forked dsDNA sequence used for capturing the DruE–dsDNA complex. The highlighted nucleotides are built into the cryo-EM model, which fits the density best. The solid lines in the sequence represent base pairing, and the dotted lines represent unwound base pairs.

(E) Close-up view of partially unwound dsDNA observed in the complex, with the cryo-EM map overlaid.

(F) Residue conservation of F1327, P1328, and W1480, which interact with the nucleobase of the tracking strand.

(G) Detailed view of the interaction between DruE residue W1480 and the DNA tracking strand.

(H) Detailed view of the interactions between DruE residues F1327 and P1328 and the DNA tracking strand.

(I) Structural comparison of DruE protomer A with protomer B in the DruE dsDNA complex.

(J) Same as in (G) but rotated 180° to show that the DNA tracking strand and displaced strand translocate through different paths.

**Figure S5. DruH DNA binding characterization**

(A) Druantia system proteins co-occurrence network.

(B) Mass-photometry characterization of DruH.

(C) Mass-photometry characterization of the interaction between DruH and DruE.

(D) Mass-photometry characterization of DruH in the presence or absence of ssDNA.

(E) Mass-photometry characterization of the interaction between DruH and DruE in the presence of ssDNA.

(F) Mass-photometry characterization of the interaction between DruH, DruE, forked dsDNA in the presence or absence of ATP. Data in (B)-(F) shown represent combined distributions from three independent measurements.

(G) DruH DNA binding with difference ssDNA substrates (**Table S2**).

(H) DruH ssDNA binding in the absence of ions or in the presence of EDTA, Zn^2+^, Mg^2+^, Ca^2+^, Co^2+^.

(I) TEB–urea denaturing gel with the same samples as in (H).

**Figure S6. Cryo-EM dataset processing results of DruH apo, in complex with ssDNA,**

(A) Representative 2D classes of the DruH apo form, the cryo-EM density map colored by local resolution (Å) estimated in cryoSPARC using the gold-standard criterion (0.143), and Fourier shell correlation (GSFSC) curves of the refined map with particle angles and poses.

(B) Five DruH models predicted by AlphaFold, colored by the pLDDT confidence value.

(C) Coulombic electrostatic potential of the DruH apo state in units of kcal/(mol·e) at 298 K.

(D) Representative 2D classes of DruH in complex with ssDNA, the cryo-EM density map colored by local resolution (Å) estimated in cryoSPARC using the gold-standard criterion (0.143), and Fourier shell correlation (GSFSC) curves of the refined complex with particle angles and poses.

(E) B-factor comparison of DruH apo and DruH–ssDNA complex models, showing reduced B-factors upon ssDNA binding, including the ssDNA binding domains.

(F) Position-dependent Z-score analysis of the DruH–ssDNA complex compared with DruH apo, indicating no major conformational changes between the models.

(G) Structural comparison of DruH apo and DruH–ssDNA complex models, with the root-mean-square deviation (RMSD) values mapped onto DruH in the complex.

**Figure S7. Cryo-EM dataset processing results of DruH in complex with ATP.**

(A) DruH ssDNA binding in the presence of 1 mM NTPs, and a TEB–urea denaturing gel showing the same samples.

(B) DruH ssDNA binding in the presence of different ATP concentration (0.06 mM, 0.25 mM, 1.00 mM and 4.00 mM).

(C) Representative 2D classes of DruH in complex with ATP, the cryo-EM density map colored by local resolution (Å) estimated in cryoSPARC using the gold-standard criterion (0.143), and Fourier shell correlation (GSFSC) curves of the refined complex with particle angles and poses.

(D) Structural comparison of DruH apo and DruH–ATP complex models, with the root-mean-square deviation (RMSD) values mapped onto DruH in the complex.

(E) B-factor comparison of DruH apo and DruH–ssDNA complex models, showing overall reduced B-factors upon ATP binding.

(F) Position-dependent Z-score analysis of the DruH–ATP complex compared with DruH apo, indicating no major conformational changes between the models.

**Figure S8. Modification analysis of bacterial genomes and visualization of phage DNA during infection**

(A) 6mA modification analysis of bacterial genomes with or without *Ec*DruIII.

(B) 4mC modification analysis of bacterial genomes with or without *Ec*DruIII.

(C) 5mC modification analysis of bacterial genomes with or without *Ec*DruIII.

(D) 5hmC modification analysis of bacterial genomes with or without *Ec*DruIII.

## Supplementary Tables

**Table S1**. **Cryo-EM data collection, refinement and validation statistics. Table S2. Oligonucleotide substrates used in this study.**

Table S3. Mass Spectrometry analysis of the *E.coli* lysates with or without expression of the *Ec*DruIII proteins.

## Supplementary Videos

**Video S1 and S2. Visualization of phage DNA during infection using a ParB–*parS* labeling system.** Time-lapse microscopy Bas08-*parS* infection assays at an MOI=0.1 in an *E. coli* strain constitutively expressing ParB^P1^ translationally fused to mScarlet-I from the chromosome **Video S1** or empty vector **Video S2**.

## Main text references

Abramson, J. et al. (2024) “Accurate structure prediction of biomolecular interactions with AlphaFold 3,” Nature, 630(8016), pp. 493–500. Available at: 10.1038/s41586-024-07487-w.

Adobe Illustrator (2021) PCMag UK. Available at: https://uk.pcmag.com/illustration/9711/adobe-illustrator (Accessed: November 28, 2023).

Aframian, N. et al. (2025) “Expression level of anti-phage defence systems controls a trade-off between protection range and autoimmunity,” Nature Microbiology, pp. 1–9. Available at: 10.1038/s41564-025-02063-y.

Aframian, N. and Eldar, A. (2023) “Abortive infection antiphage defense systems: separating mechanism and phenotype,” Trends in Microbiology, 31(10), pp. 1003–1012. Available at: 10.1016/j.tim.2023.05.002.

Antine, S.P. et al. (2024) “Structural basis of Gabija anti-phage defence and viral immune evasion,” Nature, 625(7994), pp. 360–365. Available at: 10.1038/s41586-023-06855-2.

Aravind, L. et al. (2005) “The many faces of the helix-turn-helix domain: Transcription regulation and beyond,” FEMS Microbiology Reviews, 29(2), pp. 231–262. Available at: 10.1016/j.femsre.2004.12.008.

Batth, T.S. et al. (2019) “Protein Aggregation Capture on Microparticles Enables Multipurpose Proteomics Sample Preparation*,” Molecular & Cellular Proteomics, 18(5), pp. 1027–1035. Available at: 10.1074/mcp.TIR118.001270.

Bell, R.T. et al. (2025) “YprA family helicases provide the missing link between diverse prokaryotic immune systems.” bioRxiv, p. 2025.09.15.676423. Available at: 10.1101/2025.09.15.676423.

Buckstein, M.H., He, J. and Rubin, H. (2008) “Characterization of Nucleotide Pools as a Function of Physiological State in Escherichia coli,” Journal of Bacteriology, 190(2), pp. 718–726. Available at: 10.1128/jb.01020-07.

Burby, P.E. et al. (2018) “Discovery of a dual protease mechanism that promotes DNA damage checkpoint recovery,” PLOS Genetics, 14(7), p. e1007512. Available at: 10.1371/journal.pgen.1007512.

Capella-Gutiérrez, S., Silla-Martínez, J.M. and Gabaldón, T. (2009) “trimAl: a tool for automated alignment trimming in large-scale phylogenetic analyses,” Bioinformatics, 25(15), pp. 1972–1973. Available at: 10.1093/bioinformatics/btp348.

Casjens, S.R. and Gilcrease, E.B. (2009) “Determining DNA packaging strategy by analysis of the termini of the chromosomes in tailed-bacteriophage virions,” Methods in Molecular Biology, 502, pp. 91–111. Available at: 10.1007/978-1-60327-565-1_7.

Cheng, R. et al. (2023) “Prokaryotic Gabija complex senses and executes nucleotide depletion and DNA cleavage for antiviral defense,” Cell Host & Microbe, 31(8), pp. 1331–1344.e5. Available at: 10.1016/j.chom.2023.06.014.

Couturier, A. et al. (2023) “Real-time visualisation of the intracellular dynamics of conjugative plasmid transfer,” Nature Communications, 14(1), p. 294. Available at: 10.1038/s41467-023-35978-3.

Doron, S. et al. (2018) “Systematic discovery of antiphage defense systems in the microbial pangenome,” Science, 359(6379). Available at: 10.1126/science.aar4120.

Duncan-Lowey, B. et al. (2023) “Cryo-EM structure of the RADAR supramolecular anti-phage defense complex,” Cell, 186(5), pp. 987–998.e15. Available at: 10.1016/j.cell.2023.01.012.

Eddy, S.R. (2011) “Accelerated Profile HMM Searches,” PLOS Computational Biology, 7(10), p. e1002195. Available at: 10.1371/journal.pcbi.1002195.

Emsley, P. and Cowtan, K. (2004) “Coot: model-building tools for molecular graphics,” Acta Crystallographica. Section D, Biological Crystallography, 60(Pt 12 Pt 1), pp. 2126–2132. Available at: 10.1107/S0907444904019158.

Esvelt, K.M. et al. (2013) “Orthogonal Cas9 proteins for RNA-guided gene regulation and editing,” Nature Methods, 10(11), pp. 1116–1121. Available at: 10.1038/nmeth.2681.

Gao, Y. et al. (2023) “Molecular basis of RADAR anti-phage supramolecular assemblies,” Cell, 186(5), pp. 999–1012.e20. Available at: 10.1016/j.cell.2023.01.026.

Georjon, H. and Bernheim, A. (2023) “The highly diverse antiphage defence systems of bacteria,” Nature Reviews Microbiology [Preprint]. Available at: 10.1038/s41579-023-00934-x.

Gottlin, E.B. et al. (1998) “Catalytic mechanism of the phospholipase D superfamily proceeds via a covalent phosphohistidine intermediate,” Proceedings of the National Academy of Sciences of the United States of America, 95(16), pp. 9202–9207. Available at: 10.1073/pnas.95.16.9202.

Hampton, H.G., Watson, B.N.J. and Fineran, P.C. (2020) “The arms race between bacteria and their phage foes,” Nature, 577(7790), pp. 327–336. Available at: 10.1038/s41586-019-1894-8.

Hochhauser, D. and Sorek, R. (2025) “Manipulation of the nucleotide pool in human, bacterial and plant immunity,” Nature Reviews Immunology, pp. 1–16. Available at: 10.1038/s41577-025-01206-w.

Hoelz, D.J., Hickey, R.J. and Malkas, L.H. (2004) “DNA Replication: Prokaryotes and Yeast,” in Wiley, Encyclopedia of Life Sciences. 1st ed. Wiley. Available at: 10.1038/npg.els.0003744.

Hu, H. et al. (2023) “Ion selectivity and rotor coupling of the Vibrio flagellar sodium-driven stator unit,” Nature Communications, 14(1), p. 4411. Available at: 10.1038/s41467-023-39899-z.

Hu, H. et al. (2025) “Structure and mechanism of the Zorya anti-phage defence system,” Nature, 639(8056), pp. 1093–1101. Available at: 10.1038/s41586-024-08493-8.

Humolli, D. et al. (2025) “Completing the BASEL phage collection to unlock hidden diversity for systematic exploration of phage-host interactions.” bioRxiv, p. 2024.09.12.612763. Available at: 10.1101/2024.09.12.612763.

Katoh, K. and Standley, D.M. (2013) “MAFFT multiple sequence alignment software version 7: improvements in performance and usability,” Molecular Biology and Evolution, 30(4), pp. 772–780. Available at: 10.1093/molbev/mst010.

Ledvina, H.E. and Whiteley, A.T. (2024) “Conservation and similarity of bacterial and eukaryotic innate immunity,” Nature Reviews Microbiology, 22(7), pp. 420–434. Available at: 10.1038/s41579-024-01017-1.

Li, H. et al. (2025) “The co-localizing Zorya II, Druantia III, and ARMADA II defense systems on O-island 172 confer synergistic anti-phage defense in enterohemorrhagic Escherichia coli.” bioRxiv, p. 2025.09.22.677730. Available at: 10.1101/2025.09.22.677730.

Li, J. et al. (2024) “Structures and activation mechanism of the Gabija anti-phage system,” Nature, 629(8011), pp. 467–473. Available at: 10.1038/s41586-024-07270-x.

Li, Y. and Austin, S. (2002) “The P1 plasmid is segregated to daughter cells by a ‘capture and ejection’ mechanism coordinated with Escherichia coli cell division,” Molecular Microbiology, 46(1), pp. 63–74. Available at: 10.1046/j.1365-2958.2002.03156.x.

Liebschner, D. et al. (2019) “Macromolecular structure determination using X-rays, neutrons and electrons: recent developments in Phenix,” Acta Crystallographica Section D: Structural Biology, 75(10), pp. 861–877. Available at: 10.1107/S2059798319011471.

Liu, H. and Naismith, J.H. (2008) “An efficient one-step site-directed deletion, insertion, single and multiple-site plasmid mutagenesis protocol,” BMC Biotechnology, 8(1), p. 91. Available at: 10.1186/1472-6750-8-91.

Lugmayr, W. et al. (2023) “StarMap: a user-friendly workflow for Rosetta-driven molecular structure refinement,” Nature Protocols, 18(1), pp. 239–264. Available at: 10.1038/s41596-022-00757-9.

Maffei, E. et al. (2021) “Systematic exploration of Escherichia coli phage–host interactions with the BASEL phage collection,” PLOS Biology, 19(11), p. e3001424. Available at: 10.1371/journal.pbio.3001424.

Manthei, K.A. et al. (2024) “Structural and biochemical characterization of the mitomycin C repair exonuclease MrfB,” Nucleic Acids Research, 52(11), pp. 6347–6359. Available at: 10.1093/nar/gkae308.

Mariano, G. et al. (2025) “Modularity of Zorya defense systems during phage inhibition,” Nature Communications, 16(1), p. 2344. Available at: 10.1038/s41467-025-57397-2.

Mayo-Muñoz, D. et al. (2024) “Inhibitors of bacterial immune systems: discovery, mechanisms and applications,” Nature Reviews Genetics, 25(4), pp. 237–254. Available at: 10.1038/s41576-023-00676-9.

Molineux, I.J. and Panja, D. (2013) “Popping the cork: mechanisms of phage genome ejection,” Nature Reviews Microbiology, 11(3), pp. 194–204. Available at: 10.1038/nrmicro2988.

Ofir, G. et al. (2018) “DISARM is a widespread bacterial defence system with broad anti-phage activities,” Nature Microbiology, 3(1), pp. 90–98. Available at: 10.1038/s41564-017-0051-0.

Ojima, S. et al. (2024) “Systematic Discovery of Phage Genes that Inactivate Bacterial Immune Systems.” bioRxiv, p. 2024.04.14.589459. Available at: 10.1101/2024.04.14.589459.

Payne, L.J. et al. (2022) “PADLOC: a web server for the identification of antiviral defence systems in microbial genomes,” Nucleic Acids Research, 50(W1), pp. W541–W550. Available at: 10.1093/nar/gkac400.

Perez-Riverol, Y. et al. (2025) “The PRIDE database at 20 years: 2025 update,” Nucleic Acids Research, 53(D1), pp. D543–D553. Available at: 10.1093/nar/gkae1011.

Perkins, D.N. et al. (1999) “Probability-based protein identification by searching sequence databases using mass spectrometry data,” Electrophoresis, 20(18), pp. 3551–3567. Available at: 10.1002/(SICI)1522-2683(19991201)20:18%3C3551::AID-ELPS3551%3E3.0.CO;2-2.

Pettersen, E.F. et al. (2021) “UCSF ChimeraX: Structure visualization for researchers, educators, and developers,” Protein Science: A Publication of the Protein Society, 30(1), pp. 70–82. Available at: 10.1002/pro.3943.

Punjani, A. et al. (2017) “cryoSPARC: algorithms for rapid unsupervised cryo-EM structure determination,” Nature Methods, 14(3), pp. 290–296. Available at: 10.1038/nmeth.4169.

Roske, J.J. et al. (2021) “A skipping rope translocation mechanism in a widespread family of DNA repair helicases,” Nucleic Acids Research, 49(1), pp. 504–518. Available at: 10.1093/nar/gkaa1174.

Schmidt, A. et al. (2016) “The quantitative and condition-dependent Escherichia coli proteome,” Nature Biotechnology, 34(1), pp. 104–110. Available at: 10.1038/nbt.3418.

Sievers, F. and Higgins, D.G. (2018) “Clustal Omega for making accurate alignments of many protein sequences,” Protein Science: A Publication of the Protein Society, 27(1), pp. 135–145. Available at: 10.1002/pro.3290.

Steinegger, M. and Söding, J. (2017) “MMseqs2 enables sensitive protein sequence searching for the analysis of massive data sets,” Nature Biotechnology, 35(11), pp. 1026–1028. Available at: 10.1038/nbt.3988.

Taylor, J.A. et al. (2015) “Specific and non-specific interactions of ParB with DNA: implications for chromosome segregation,” Nucleic Acids Research, 43(2), pp. 719–731. Available at: 10.1093/nar/gku1295.

Tesson, F., Planel, Remi, et al. (2024) “A Comprehensive Resource for Exploring Antiphage Defense: DefenseFinder Webservice, Wiki and Databases.” bioRxiv, p. 2024.01.25.577194. Available at: 10.1101/2024.01.25.577194.

Tesson, F., Planel, Rémi, et al. (2024) “A Comprehensive Resource for Exploring Antiphage Defense: DefenseFinder Webservice,Wiki and Databases,” Peer Community Journal, 4. Available at: 10.24072/pcjournal.470.

Tesson, F. and Bernheim, A. (2023) “Synergy and regulation of antiphage systems: toward the existence of a bacterial immune system?,” Current Opinion in Microbiology, 71, p. 102238. Available at: 10.1016/j.mib.2022.102238.

Tyanova, S. et al. (2016) “The Perseus computational platform for comprehensive analysis of (prote)omics data,” Nature Methods, 13(9), pp. 731–740. Available at: 10.1038/nmeth.3901.

Volkmer, B. and Heinemann, M. (2011) “Condition-Dependent Cell Volume and Concentration of Escherichia coli to Facilitate Data Conversion for Systems Biology Modeling,” PLOS ONE, 6(7), p. e23126. Available at: 10.1371/journal.pone.0023126.

Wang, S. et al. (2023) “Landscape of New Nuclease-Containing Antiphage Systems in Escherichia coli and the Counterdefense Roles of Bacteriophage T4 Genome Modifications,” Journal of Virology, 97(6), pp. e00599–23. Available at: 10.1128/jvi.00599-23.

Wong, T.K.F. et al. (2025) “IQ-TREE 3: Phylogenomic Inference Software using Complex Evolutionary Models.” Available at: https://ecoevorxiv.org/repository/view/8916/ (Accessed: February 21, 2026).

Wu, Y. et al. (2024) “Bacterial defense systems exhibit synergistic anti-phage activity,” Cell Host & Microbe [Preprint]. Available at: 10.1016/j.chom.2024.01.015.

Xu, Z. et al. (2026) “The Ppl protein senses 3′-hydroxyl DNA overhangs and NTP depletion to halt phage infection,” Molecular Cell, 86(1), pp. 180–193.e6. Available at: 10.1016/j.molcel.2025.11.011.

Yang, W. (2011) “Nucleases: diversity of structure, function and mechanism,” Quarterly Reviews of Biophysics, 44(1), pp. 1–93. Available at: 10.1017/S0033583510000181.

Youngren, B. et al. (2014) “The multifork Escherichia coli chromosome is a self-duplicating and self-segregating thermodynamic ring polymer,” Genes & Development, 28(1), pp. 71–84. Available at: 10.1101/gad.231050.113.

Zhang, H. et al. (2025) “The two-component nuclease-active KELShedu system confers broad antiphage activity via abortive infection,” Science Advances, 11(44), p. eadv4747. Available at: 10.1126/sciadv.adv4747.

