## Supplemental data for "Molecular basis of the Druantia anti-phage defense system"

### Inventory:

**Supplementary Figures 1-8**

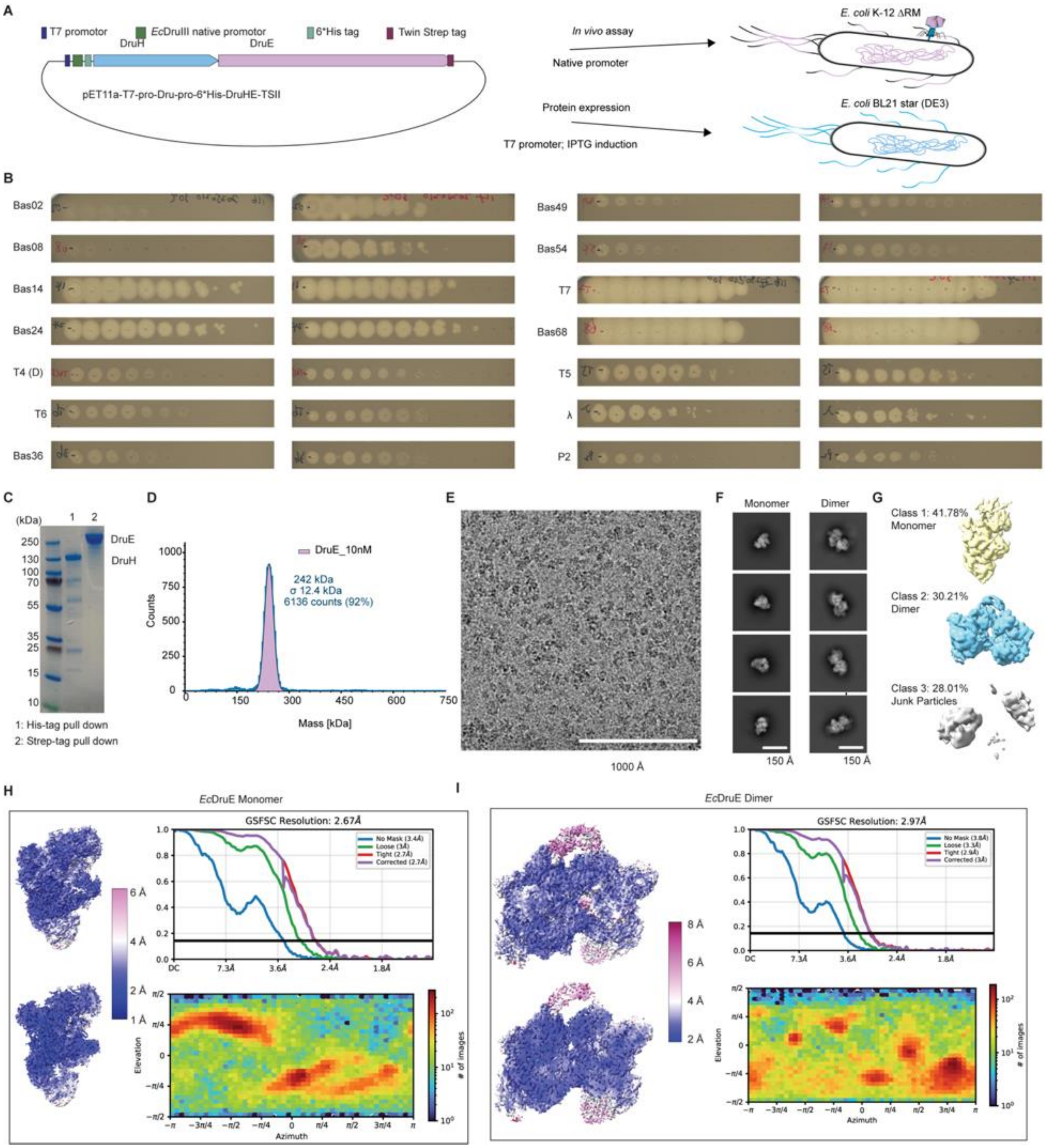


**Figure S1. DruE exits as monomer and dimer and cryo-EM dataset processing results**

(A) Schematic representation of the cloning strategy for *Ec*DruIII.

(B) Plaque assays of phages challenging *E. coli* K-12 ΔRM expressing either a vector control or *Ec*DruIII. Images are representative of three replicates.

(C) Representative SDS-PAGE gel of the co-purifying DruH and DruE proteins. The first lane shows co-purification using a His-tag on DruH, and the second lane shows co-purification using a twin Strep-tag on DruE. Neither affinity-purified DruH nor affinity-purified DruE pulled down the other protein. Images are representative of three replicates.

(D) Mass photometry analysis of purified DruE at 10 nM concentration. The data shown represent combined distributions from three independent measurements.

(E) Cryo-EM image of the DruE sample under cryogenic conditions.

(F) Representative 2D class average of the DruE monomer and dimer.

(G) 3D classes of DruE apo dataset with particle populations.

(H) Cryo-EM density map of the DruE monomer colored by local resolution (Å), estimated in cryoSPARC, and gold-standard (0.143) Fourier shell correlation (GSFSC) curves of the refined complex with particle angles and poses.

(I) Cryo-EM density map of the DruE dimer colored by local resolution (Å), estimated in cryoSPARC and gold-standard (0.143) Fourier shell correlation (GSFSC) curves of the refined EM map with particle angles and poses.

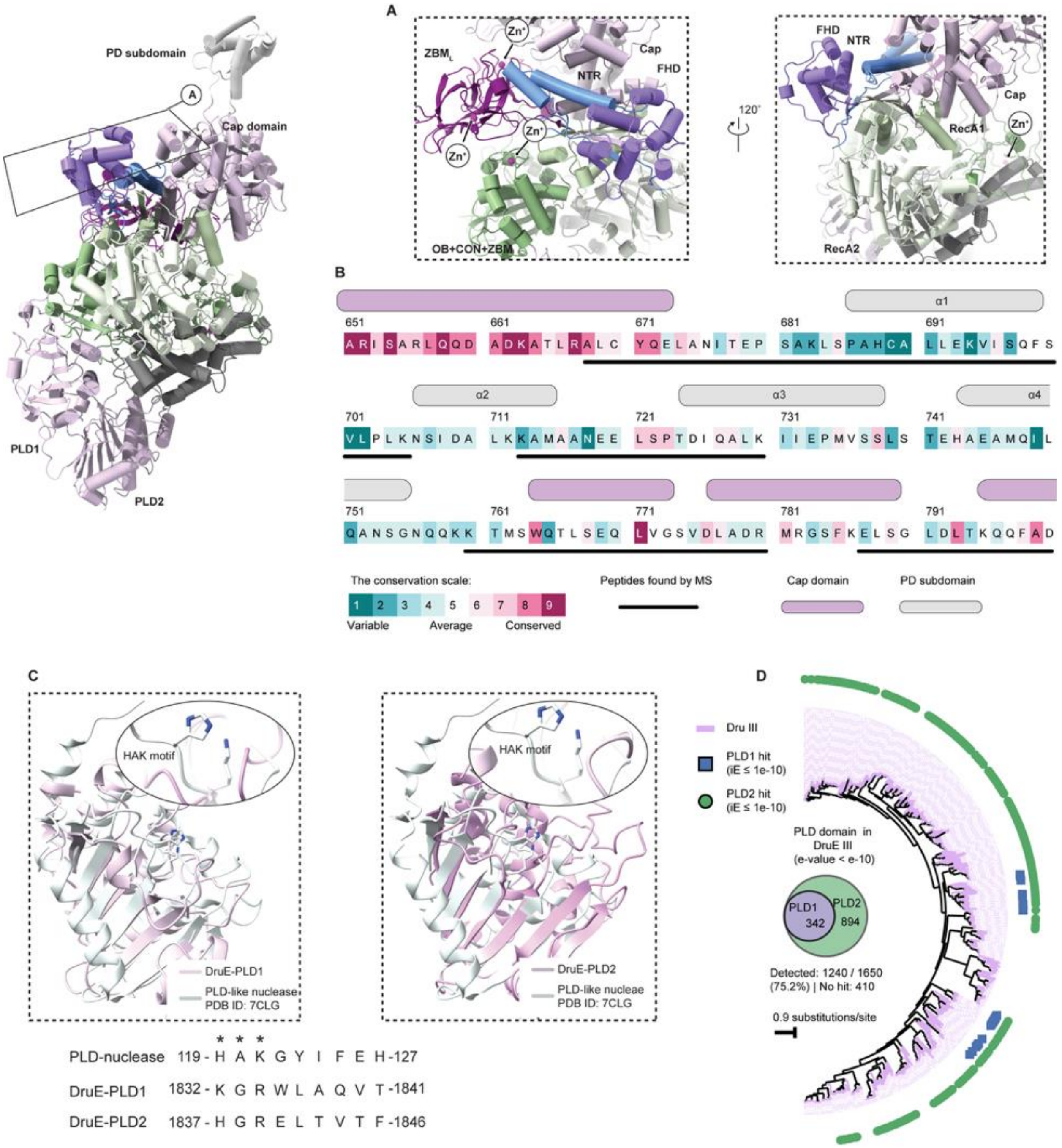


**Figure S2. DruE structure and domains**

(A) Zoom-in view of the DruE structute, showing that the DruE NTR spatially locks the core folds.

(B) Amino acids and secondary structure of the DruE PD subdomain with part of the Cap domain. Peptides identified by mass spectrometry that cover the DruE PD subdomain are indicated as black lines beneath the residues.

(C) Superimposition of DruE PLD1 and PLD2 with the top hit (PDB ID: 7CLG) from Foldseek.

(D) Phylogeny of DruIIIE sequences, with the presence of PLD1 and/or PLD2 subdomain indicated.

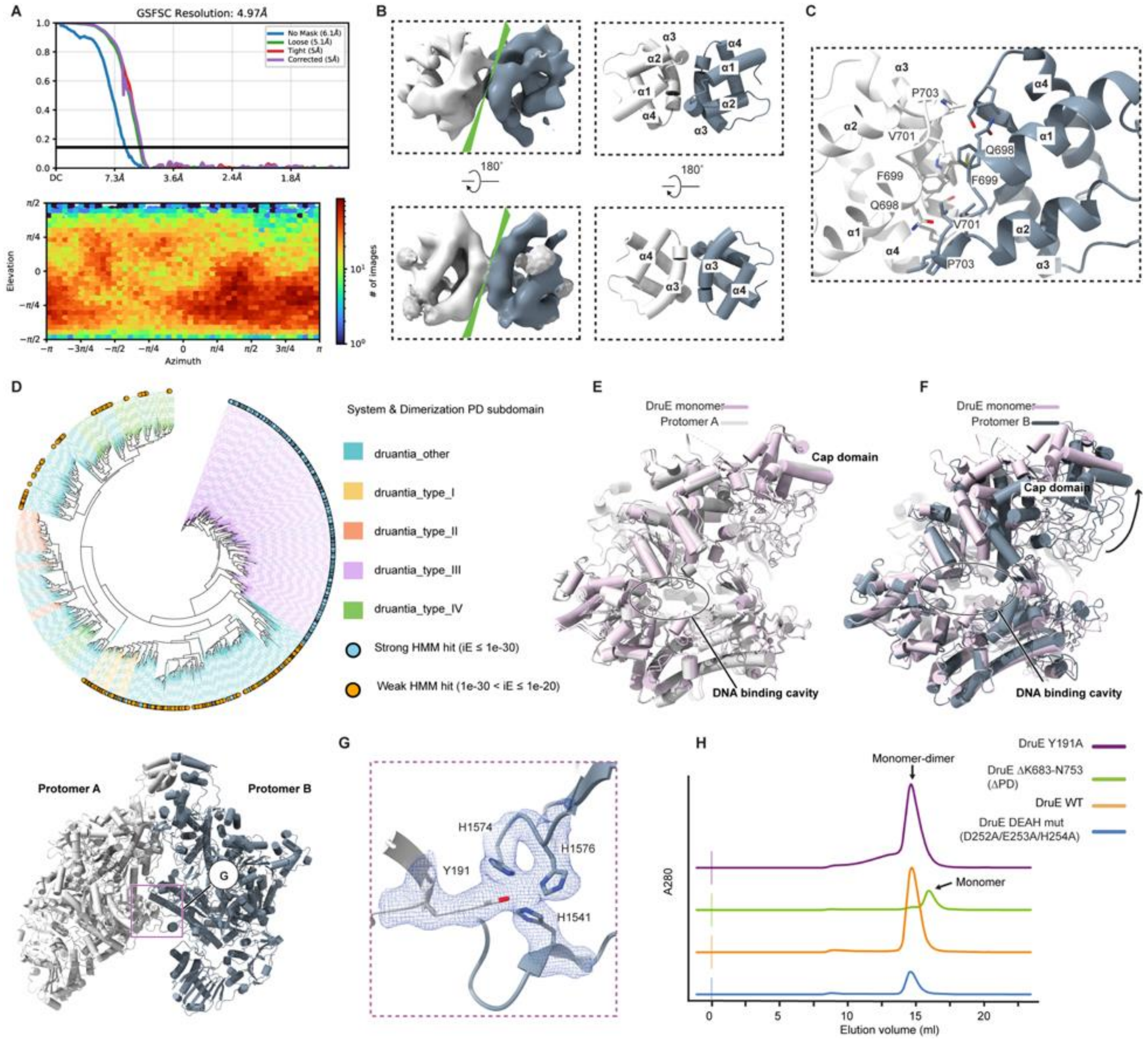


**Figure S3. Structure details of DruE PD subdomains, core domains**

(A) Gold-standard (0.143) Fourier shell correlation (GSFSC) curves of the refined DruE PD subdomain from the DruE dimer, with particle angles and poses.

(B) Cryo-EM density map and model of the dimerized DruE PD subdomain.

(C) Zoomed-in view of the dimerized PD subdomain interface.

(D) Phylogeny of DruE sequences, with HMM hits for the dimerized PD subdomain indicated across Durantia families.

(E) Structural comparison of the DruE monomer with protomer A in the DruE asymmetric dimer.

(F) Structural comparison of the DruE monomer with protomer B in the DruE asymmetric dimer. In (E) and (F), the PD subdomain, PLD1, and PLD2 are excluded.

(G) Close-up view of the interactions at domain assembly interface III, mediated by Y191 from protomer A and three histidines from protomer B, with the cryo-EM map overlaid.

(H) Size-exclusion chromatography (SEC) of the purified Y191A mutant in comparison with DruE WT, DruE ΔK683-N753 (ΔPD) and DruE DEAH mutant (D252A/E253A/H254A).

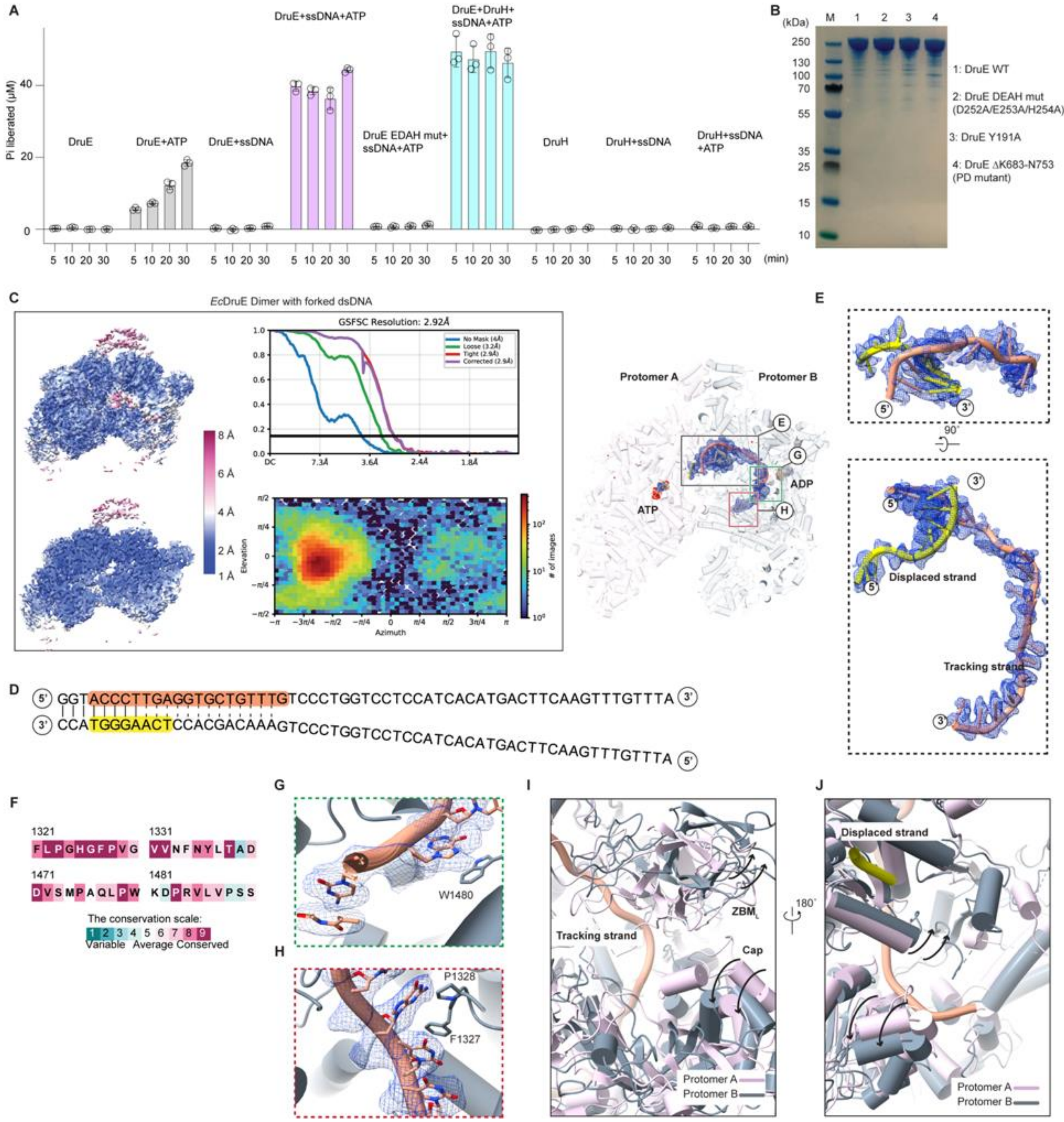


**Figure S4. Structure of DruE in complex with forked dsDNA**

(A) Malachite green ATPase assay of DruE, DruE mutant or DruH with or without ssDNA.

(B) Representative SDS-PAGE gel of the purified DruE proteins used for helicase assays. Images are representative of at least three replicates.

(C) Cryo-EM density map of the DruE dimer in complex with forked dsDNA, colored by local resolution (Å) estimated in cryoSPARC, and gold-standard (0.143) Fourier shell correlation (GSFSC) curves of the refined complex with particle angles and poses.

(D) Forked dsDNA sequence used for capturing the DruE–dsDNA complex. The highlighted nucleotides are built into the cryo-EM model, which fits the density best. The solid lines in the sequence represent base pairing, and the dotted lines represent unwound base pairs.

(E) Close-up view of partially unwound dsDNA observed in the complex, with the cryo-EM map overlaid.

(F) Residue conservation of F1327, P1328, and W1480, which interact with the nucleobase of the tracking strand.

(G) Detailed view of the interaction between DruE residue W1480 and the DNA tracking strand.

(H) Detailed view of the interactions between DruE residues F1327 and P1328 and the DNA tracking strand.

(I) Structural comparison of DruE protomer A with protomer B in the DruE dsDNA complex.

(J) Same as in (G) but rotated 180° to show that the DNA tracking strand and displaced strand translocate through different paths.

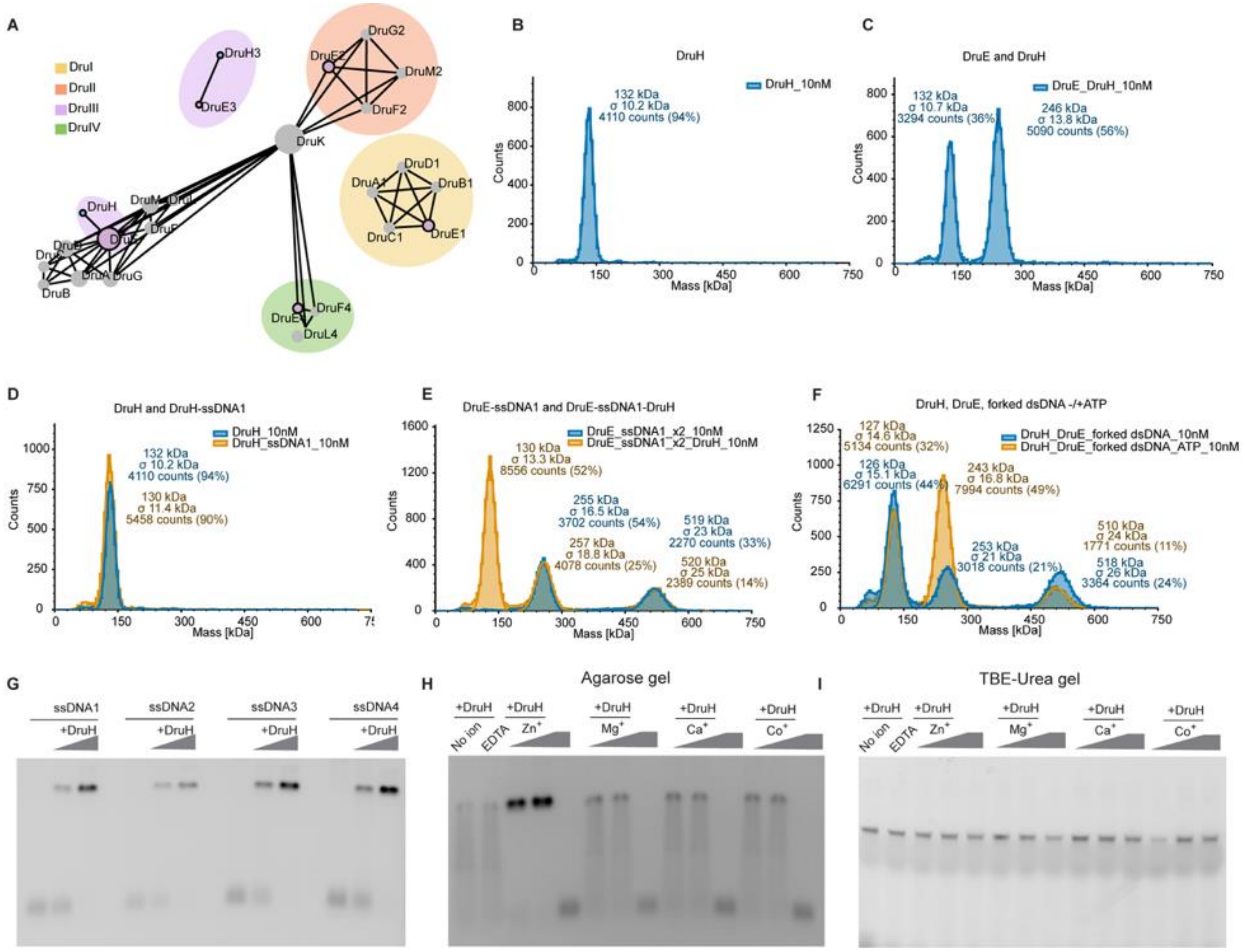


**Figure S5. DruH DNA binding characterization**

(A) Druantia system proteins co-occurrence network.

(B) Mass-photometry characterization of DruH.

(C) Mass-photometry characterization of the interaction between DruH and DruE.

(D) Mass-photometry characterization of DruH in the presence or absence of ssDNA.

(E) Mass-photometry characterization of the interaction between DruH and DruE in the presence of ssDNA.

(F) Mass-photometry characterization of the interaction between DruH, DruE, forked dsDNA in the presence or absence of ATP. Data in (B)-(F) shown represent combined distributions from three independent measurements.

(G) DruH DNA binding with difference ssDNA substrates (**Table S2**).

(H) DruH ssDNA binding in the absence of ions or in the presence of EDTA, $Zn^{2+}$, $Mg^{2+}$, $Ca^{2+}$, $Co^{2+}$.

(I) TEB–urea denaturing gel with the same samples as in (H).

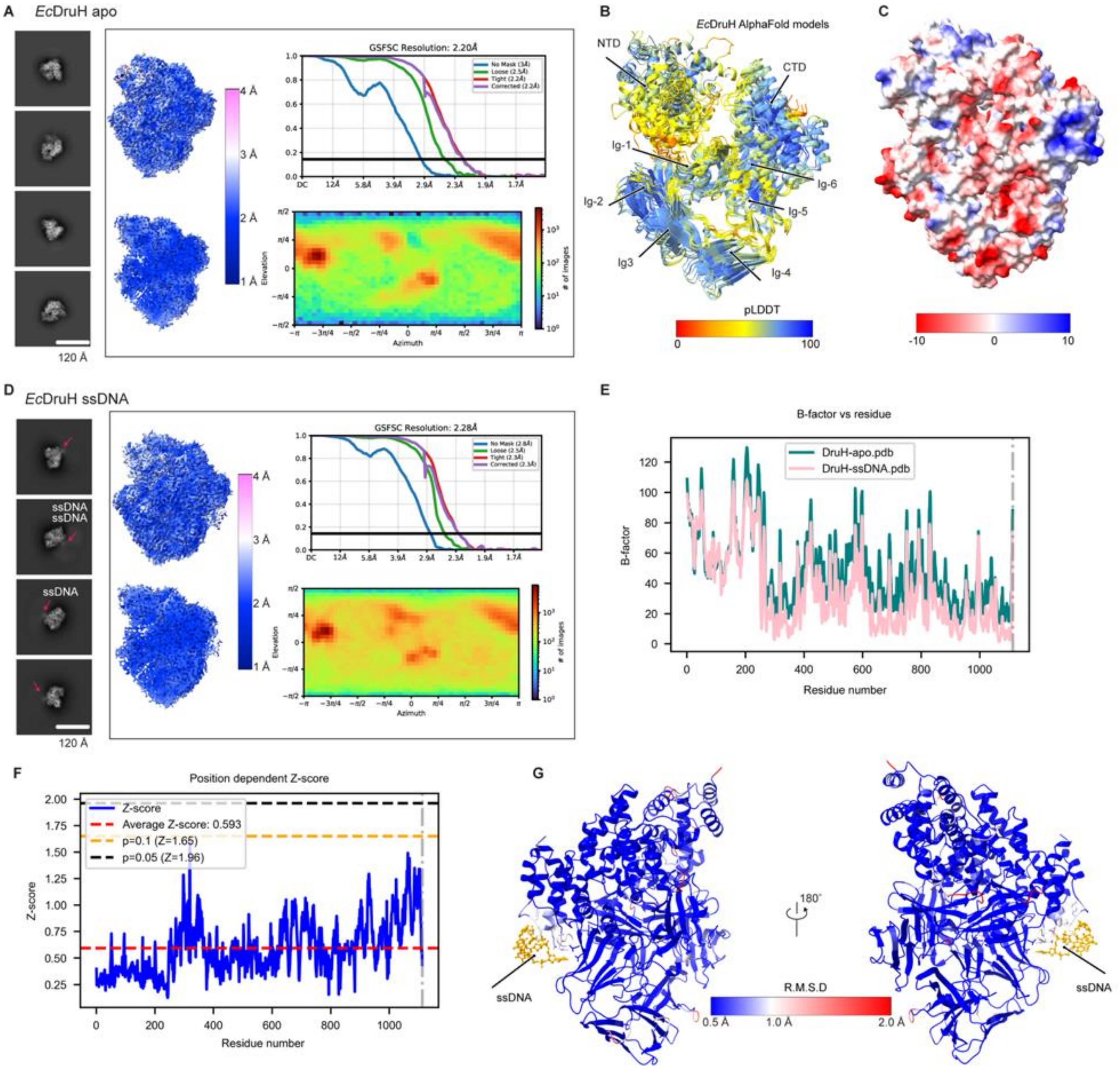


**Figure S6. Cryo-EM dataset processing results of DruH apo, in complex with ssDNA**

(A) Representative 2D classes of the DruH apo form, the cryo-EM density map colored by local resolution (Å) estimated in cryoSPARC using the gold-standard criterion (0.143), and Fourier shell correlation (GSFSC) curves of the refined map with particle angles and poses.

(B) Five DruH models predicted by AlphaFold, colored by the pLDDT confidence value.

(C) Coulombic electrostatic potential of the DruH apo state in units of kcal/(mol·e) at 298 K.

(D) Representative 2D classes of DruH in complex with ssDNA, the cryo-EM density map colored by local resolution (Å) estimated in cryoSPARC using the gold-standard criterion (0.143), and Fourier shell correlation (GSFSC) curves of the refined complex with particle angles and poses.

(E) B-factor comparison of DruH apo and DruH–ssDNA complex models, showing reduced B-factors upon ssDNA binding, including the ssDNA binding domains.

(F) Position-dependent Z-score analysis of the DruH–ssDNA complex compared with DruH apo, indicating no major conformational changes between the models.

(G) Structural comparison of DruH apo and DruH–ssDNA complex models, with the root-mean-square deviation (RMSD) values mapped onto DruH in the complex.

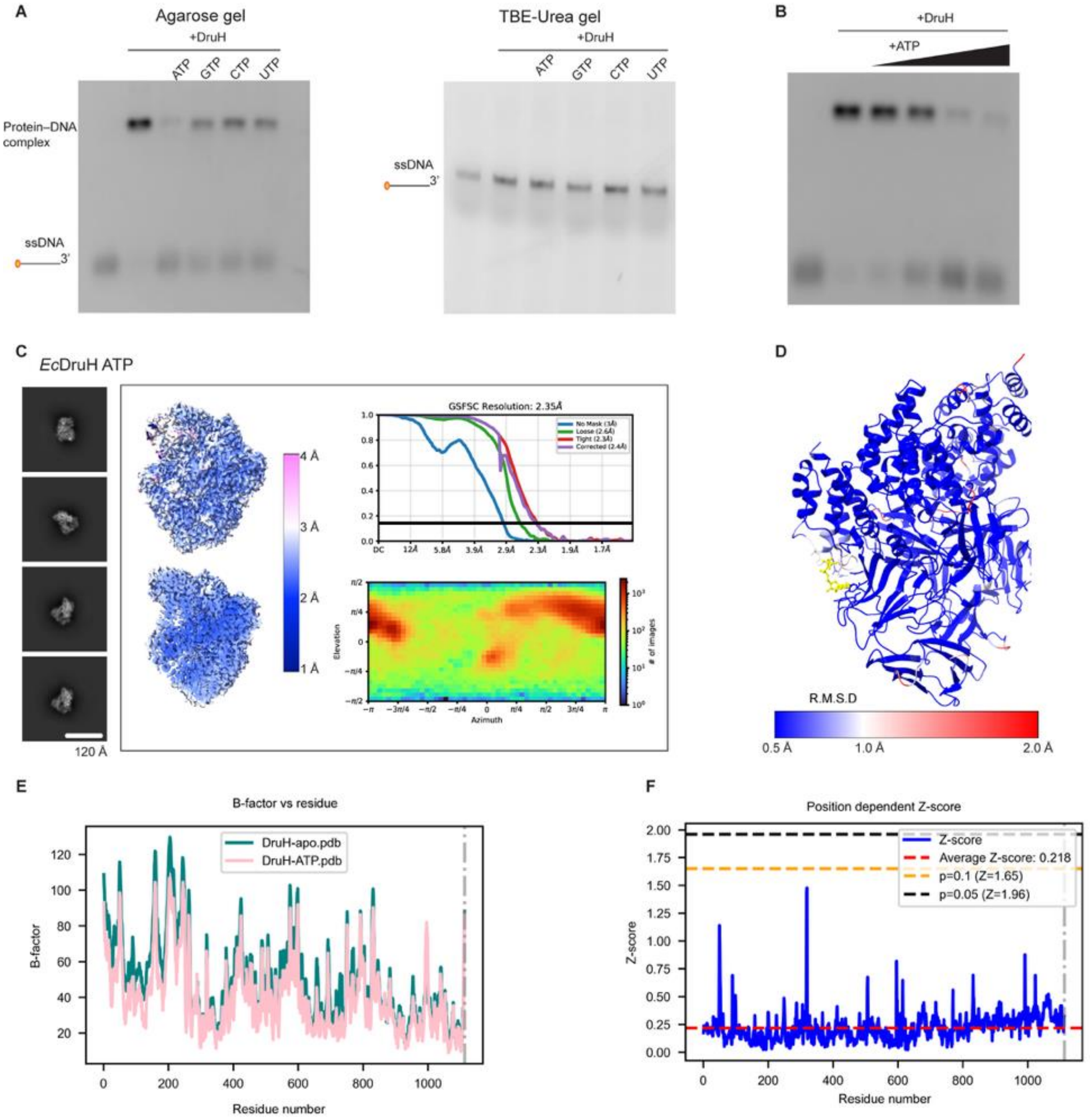


**Figure S7. Cryo-EM dataset processing results of DruH in complex with ATP.**

(A) DruH ssDNA binding in the presence of 1 mM NTPs, and a TEB–urea denaturing gel showing the same samples.
(B) DruH ssDNA binding in the presence of different ATP concentration (0.06 mM, 0.25 mM, 1.00 mM and 4.00 mM).
(C) Representative 2D classes of DruH in complex with ATP, the cryo-EM density map colored by local resolution (Å) estimated in cryoSPARC using the gold-standard criterion (0.143), and Fourier shell correlation (GSFSC) curves of the refined complex with particle angles and poses.

(D) Structural comparison of DruH apo and DruH–ATP complex models, with the root-mean-square deviation (RMSD) values mapped onto DruH in the complex.

(E) B-factor comparison of DruH apo and DruH–ssDNA complex models, showing overall reduced B-factors upon ATP binding.

(F) Position-dependent Z-score analysis of the DruH–ATP complex compared with DruH apo, indicating no major conformational changes between the models.

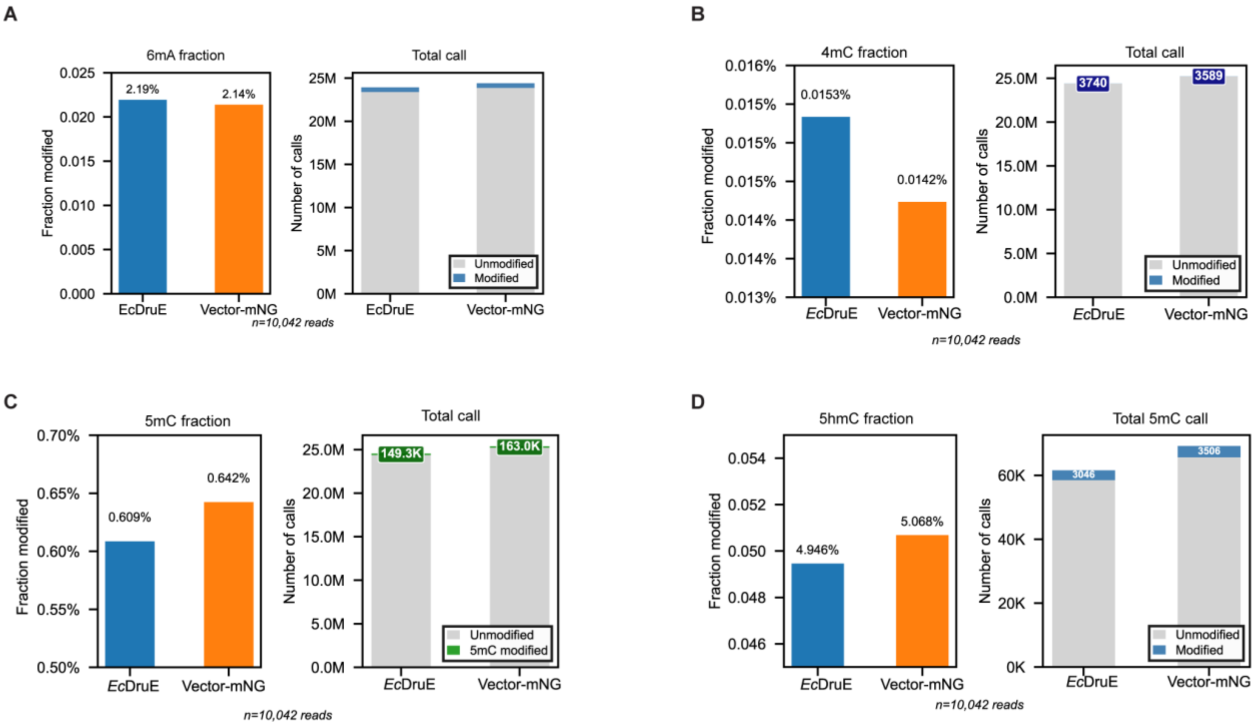


**Figure S8. Modification analysis of bacterial genomes**

(A) 6mA modification analysis of bacterial genomes with or without *Ec*DruIII.

(B) 4mC modification analysis of bacterial genomes with or without *Ec*DruIII.

(C) 5mC modification analysis of bacterial genomes with or without *Ec*DruIII.

(D) 5hmC modification analysis of bacterial genomes with or without *Ec*DruIII.
